# Inhibition of the glucocorticoid-activating enzyme 11β-hydroxysteroid dehydrogenase type 1 drives concurrent 11-oxygenated androgen excess

**DOI:** 10.1101/2023.06.05.543687

**Authors:** Lina Schiffer, Imken Oestlund, Jacky Snoep, Lorna C. Gilligan, Angela E. Taylor, Alexandra J. Sinclair, Rishi Singhal, Adrian Freeman, Ramzi Ajjan, Ana Tiganescu, Wiebke Arlt, Karl-Heinz Storbeck

**Author notes:** Shared first authors (Lina Schiffer, Imken Oestlund). **Correspondence:** Karl-Heinz Storbeck, Department of Biochemistry, Stellenbosch University, Private Bag X1, Matieland, 7602, South Africa.

## Abstract

Aldo-keto reductase 1C3 (AKR1C3) is a key enzyme in the activation of both classic and 11-oxygenated androgens. In adipose tissue, AKR1C3 is co-expressed with 11β-hydroxysteroid dehydrogenase type 1 (HSD11B1), which catalyses the local activation of glucocorticoids but also the inactivation of 11-oxygenated androgens, and thus has the potential to counteract AKR1C3. Using a combination of *in vitro* assays and *in silico* modelling we show that HSD11B1 attenuates the biosynthesis of the potent 11-oxygenated androgen, 11-ketotestosterone, by AKR1C3. Employing *ex vivo* incubations of human female adipose tissue samples we show that inhibition of HSD11B1 results in the increased peripheral biosynthesis of 11-ketotestosterone. Moreover, circulating 11KT increased 2-3 fold in individuals with type 2 diabetes after receiving the selective oral HSD11B1 inhibitor AZD4017 for 35 days, thus confirming that HSD11B1 inhibition results in systemic increases in 11KT concentrations. Our findings show that HSD11B1 protects against excess 11KT production by adipose tissue, a finding of particular significance when considering the evidence for adverse metabolic effects of androgens in women. Therefore, when targeting glucocorticoid activation by HSD11B1 inhibitor treatment in women, the consequently increased generation of 11-ketotestosterone may offset beneficial effects of decreased glucocorticoid activation.

**Graphical Abstract:** 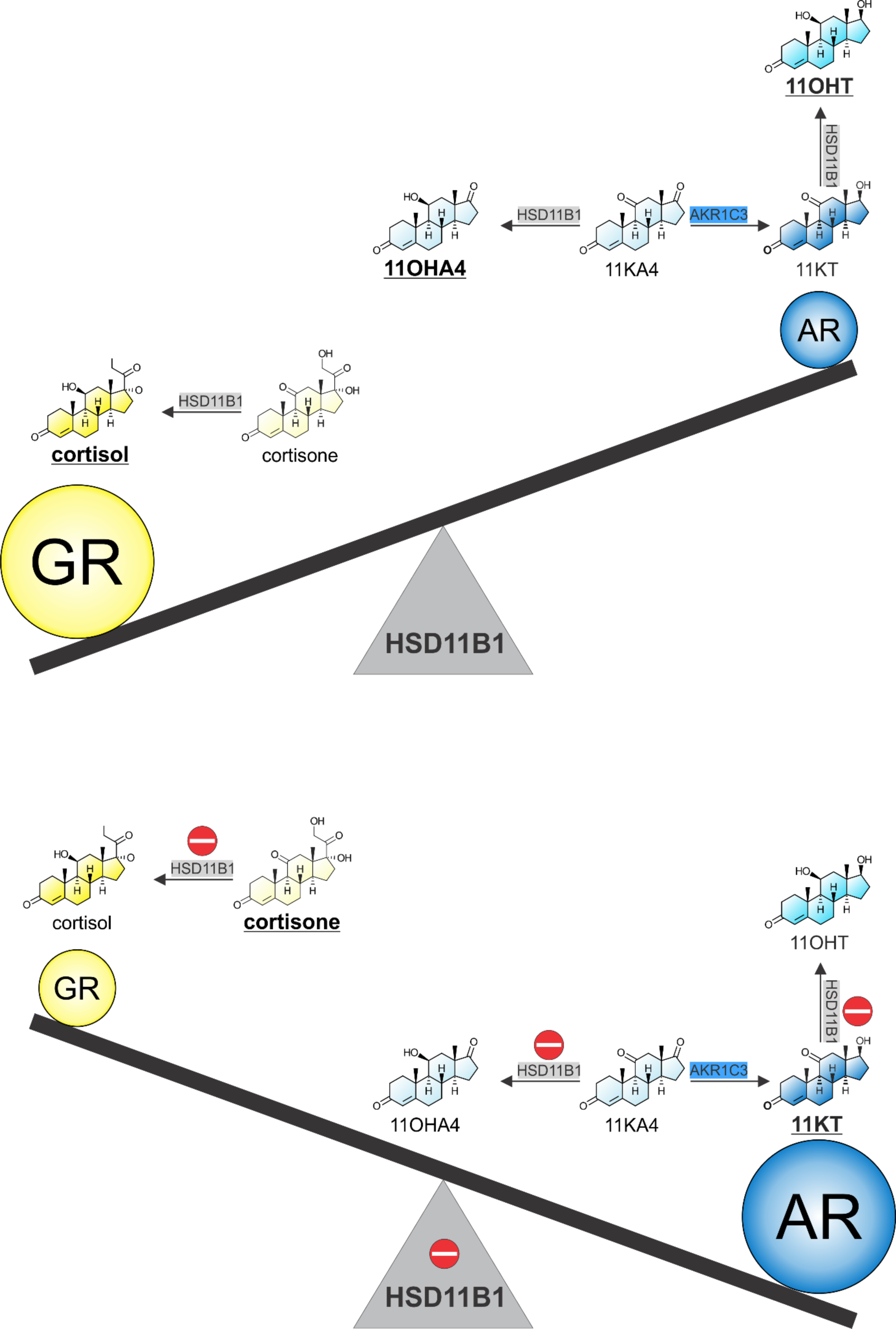

## Introduction

The 11-oxygenated androgens are a group of adrenal-derived and peripherally activated androgens that make a significant contribution to the circulating androgen pool, particularly in women (>>>). The most potent 11-oxygenated androgen in circulation is 11-ketotestosterone (11KT), which has been shown to bind to and activate the androgen receptor with an affinity and potency similar to testosterone (5). Moreover, several studies have also shown that 11KT circulates at concentrations equal to or higher than those of testosterone in women (3, 6, 7). Unlike the classic androgens, 11-oxygenated androgens do not decline with age, with the result that 11KT is the most abundant active androgen in the circulation of postmenopausal women (3, 6). While the role of 11-oxygenated androgens in healthy individuals has yet to be defined, these androgens have been shown to play a role in disease states such as polycystic ovary syndrome (PCOS), congenital adrenal hyperplasia (CAH), Cushing’s syndrome and castration resistant prostate cancer (CRPC) (8–12).

The biosynthesis of 11KT begins in the adrenal, where cytochrome P450 11B1 (CYP11B1) catalyses the 11β-hydroxylation of locally produced androstenedione to yield 11β-hydroxyandrostenedione (11OHA4). 11OHA4 is released into circulation, and serves as the primary precursor for the biosynthesis of 11KT by peripheral tissues (**Figure 1**). Specifically, 11OHA4 is converted to 11-ketoandrostenedione (11KA4) by mineralocorticoid target tissues expressing 11β-hydroxysteroid dehydrogenase type 2 (HSD11B2), such as the kidney (1>). 11KA4 then subsequently serves as the substrate for the key androgen activating enzyme, aldo-keto reductase 1C3 (AKR1C3), which is abundantly expressed in adipose tissue (14–17). Notably, AKR1C3 catalyses the conversion of 11KA4 to 11KT significantly more efficiently than that of androstenedione to testosterone (16, 17). The biosynthesis of 11KT is, however, more complex than that of testosterone as the presence of the 11-keto group makes both 11KA4 and 11KT substrates for 11β-hydroxysteroid dehydrogenase type 1 (HSD11B1)—an enzyme best known for amplification of glucocorticoid signalling, which it achieves by converting the inactive cortisone to active cortisol in key metabolic tissues including the liver, adipose tissue and skeletal muscle (11, 18–20). Given that AKR1C3 and HSD11B1 are co-expressed in adipose tissue, we propose that HSD11B1 regulates the amount of 11KT produced in two ways. First, by competing with AKR1C3 for 11KA4, thereby limiting the amount of 11KA4 converted to 11KT by AKR1C3. Secondly, by converting some of the 11KT formed to the far less androgenic 11β-hydroxytestosterone (11OHT) (11) (**Figure 1**). Indeed, while investigating the metabolism of 11KA4 in differentiated Simpson-Golabi-Behmel syndrome preadipocyte (SGBS) cells Paulukinas et. al. noted that HSD11B1 may be protective of androgen excess (17).

**Figure 1:**
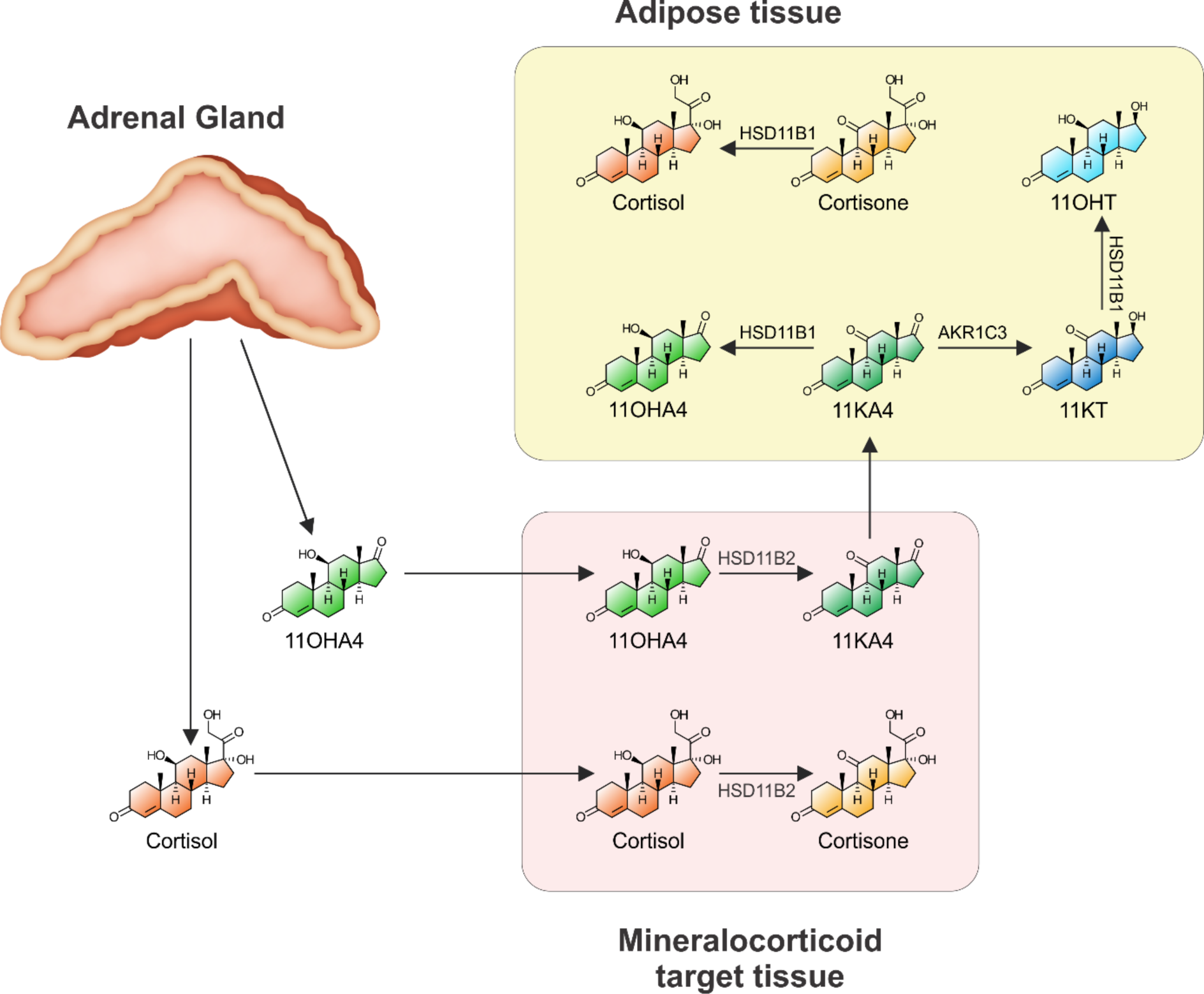
Schematic showing the peripheral metabolism of cortisol and 11OHA4. Cortisol is converted to its inactive form, cortisone, by HSD11B2 expressed in mineralocorticoid target tissues. Cortisone is in turn converted back to cortisol by HSD11B1 expressed in glucocorticoid target tissues (e.g., adipose). Like cortisol, adrenal derived 11OHA4 is a substrate for HSD11B2, yielding 11KA4. 11KA4 is a substrate for both the androgen activating enzyme AKR1C3 and HSD11B1, both of which are expressed in adipose tissue. Conversion of 11KA4 to 11OHA4 by HSD11B1 prevents the conversion of 11KA4 to the potent 11-oxygenated androgen, 11KT. The 11KT that is produced by AKR1C3 is also a substrate for HSD11B1, yielding the substantially less active androgen, 11OHT.

This proposed mechanism is important when considering that HSD11B1 has long been a therapeutic target for prevention of tissue-specific excess glucocorticoid exposure and treatment of related adverse metabolic effects (19, 21–24). Promising preclinical studies in rodents, which crucially do not produce 11-oxygenated androgens (25–30), were unable to accurately model competing HSD11B1 and AKR1C3 effects in humans. Our proposed hypothesis could provide an explanation for the more limited effect sizes observed in clinical trials with HSD11B1 inhibitors (31, 32, 41, 42, 33–40), as increased 11KT production resulting from HSD11B1 inhibition could counteract beneficial effects of reduced glucocorticoid signalling, by increasing androgen signalling. This is particularly significant in women where elevated androgens are linked to insulin resistance, type 2 diabetes, hypertension, cardiovascular disease and metabolic dysfunction associated steatotic liver disease (MASLD), in particular in the context of polycystic ovary syndrome, for which androgen excess is a defining feature (43–52).

Here, we characterised the interplay between HSD11B1 and AKR1C3 as co-regulators of 11KT biosynthesis using a combination of *in vitro*, *in silico*, and human *ex vivo* and *in vivo* approaches including selective HSD11B1 inhibitor treatment in individuals with type 2 diabetes.

## Materials and Methods

### Steroids and inhibitors

Androstenedione (4-androstene-3,17-dione), carbenoxolone ((3β,20β)-3-(3-carboxy-1-oxopropoxy)-11-oxo-olean-12-en-29-oic acid, disodium salt; CBX), cortisol (11β,17α,21-trihydroxypregn-4-ene-3,20-dione), cortisone (17α,21-dihydroxy-4-pregnene-3,11,20-trione), DHEA (5-androsten-3β-ol-17-one), DHEA sulfate (5-androsten-3β-ol-17-one-3-sulfate) and testosterone (4-androsten-17β-ol-3-one) were purchased from Sigma Aldrich. 11β-hydroxyandrostenedione (11β-hydroxy-4-androstene-3,17-dione), 11β-hydroxytestosterone (11β,17β-dihydroxyandrost-4-en-3-one), 11-ketoandrostenedione (4-androsten-3,11,17-trione) and 11-ketotestosterone (4-androsten-17β-ol-3,11-dione) were from Steraloids. The deuterated internal standards 11β-hydroxyandrostenedione-2,2,4,6,6,16,16-D7 (D7-11OHA4), 11-ketotestosterone-16,16,17-D3 (D3-11KT), androstenedione-2,2,4,6,6,16,16-D7 (D7-A4), cortisol-9,11,12,12-D4 (D4-F), DHEAS-16,16-D2 (D2-DHEAS) and testosterone-16,16,17-D3 (D3-T) were all purchased from Cambridge Isotope Laboratories. Dehydroepiandrosterone-2,2,3,4,4,6-D6 (D6-DHEA) and cortisone-2,2,4,6,6,9,12,12-D8 (D8-E) were purchased from Sigma Aldrich, while 11-ketoandostenedione-2,2,4,6,6,9,12,12,16,16-D10 (D10-11KA4) was custom synthesized by Toronto Research Chemicals. The selective HSD11B1 inhibitor, 2-[(3S)-1-[5-(cyclohexylcarbamoyl)-6-propylsulfanylpyridin-2-yl]-3-piperidyl] acetic acid (AZD-4017) was provided by AstraZeneca.

### Plasmids

The pcDNA3 vector containing AKR1C3 was provided by Prof. J. Adamski (Helmholtz Zentrum München, Germany). The pcDNA3 vectors containing HSD11B1 and hexose 6-phosphate dehydrogenase (H6PDH) were from Prof. K. Chapman (University of Edinburgh, UK). The pTAT-GRE-Elb-luc luciferase promoter reporter construct, containing two copies of the GRE was from Prof. G. Jenster (Erasmus University of Rotterdam, Netherlands), while the GR expressing plasmid (pRShGRα) was from Prof. R. Evans (Howard Hughes Medical Institure, La Jolla, USA). The pCIneo plasmid expressing the wild-type human AR was from Prof. N. Sharifi (Cleveland Clinic, Cleveland, USA). A pCIneo plasmid, containing no cDNA insert, was purchased from Promega (Wisconsin, USA).

### Cell lines

HEK293 cells were purchased from the American Type Culture Collections (ATCC) and cultured in high glucose Dulbecco’s Modified Eagle Medium (DMEM) supplemented with 10% FBS, 1.5 g/L NaHCO3 and 1% penicillin-streptomycin and maintained at 37 °C in 90% humidity and 5% CO^2^. The cells were authenticated by short-tandem repeat profiling (NorthGene) and were regularly tested for mycoplasma contamination.

### Determining kinetic parameters for HSD11B1 and computational model construction

The kinetic parameters (Km, *app* and Vmax, *app*) for HSD11B1 were determined using the methodology previously reported for AKR1C3 (1>). Briefly, HEK293 cells were seeded into 10 cm^2^ tissue culture dishes at a density of 2×10^5^ cells per mL (10 mL per dish). After a 24 hour incubation period, cells were co-transfected with 0.5 µg HSD11B1 and 0.5 µg H6PDH DNA using X-tremeGENE™ HP DNA transfection reagent (Roche) according to manufacturer’s protocol. Following a 24 hour incubation the transfected cells were re-plated into 48-well Corning® CELLBIND® plates at a density of 2×10^5^ cells per mL (300 µL per well). Cells were incubated for an additional 24 hours to allow for attachment before being treated with serum free media, containing appropriate steroid substrate (cortisone, 11KA4, or 11KT) at concentrations ranging from 0.1 µM to 10 µM. Aliquots (250 µL) for the quantification of steroid metabolism by ultra-high performance liquid chromatography tandem mass spectrometry (UHPLC-MS/MS) were collected at specific time intervals. Kinetic parameters were determined in Wolfram Mathematica (Version 13) by assuming Michaelis Menten kinetics and fitting the Km, *app* and Vmax, *app* values by minimizing the sum of the squared differences between the experimentally determined progress curves and model, weighted by the variance using the NMinimize function. The model was fitted to triplicate data from three independent experiments. The transfection efficiency of each individual experiment was taken into account by determining the initial rate of the conversion of 100 nM 11KA4 for each independent experiment. A computational model containing the determined kinetic for HSD11B1 and those previously determined for AKR1C3 was subsequently constructed. This model is comprised of ordinary differential equations (ODEs) for the metabolites, and numerical integration by making use of the NDSolve routine in Mathematica allowed for experimentally comparable metabolite profiles to be obtained. Additionally, the model was used to predict the conversion of cortisone or 11KA4 for varying HSD11B1:AKR1C3 ratios, see Supplementary material for a complete model description and simulation details.

### Determining the effect of different HSD11B1:AKR1C3 ratios on cortisone and 11KA4 metabolism

HEK293 cells were seeded into 10 cm^2^ tissue culture dishes at a density of 2×10^5^ cells per mL (10 mL per dish). After a 24 hour incubation period, dishes were co-transfected with either 0.5 µg HSD11B1 and 0.5 µg H6PDH or 0.5 µg AKR1C3 and 0.5 µg pCINeo using X-tremeGENE™ HP DNA transfection reagent (Roche). Additional dishes were transfected with 1 µg of the pCINeo (no insert) as to allow for the subsequent plating of an equal number of transfected cells at all HSD11B1:AKR1C3 ratios. After 24 hours cells were counted and combined in different ratios of HSD11B1:AKR1C3, before being re-plated into 12-well Corning® CELLBIND® surface plates at a density of 4×10^5^ cells per mL (1 mL per well). After an additional incubation period of 24 hours, the media was replaced with serum free DMEM containing 100 nM substrate (11KA4 or cortisone). Aliquots (500 µL) were collected 24 hours after the addition of substrate for analysis by UHPLC-MS/MS, as described below.

### Determining the effect of CBX on HSD11B1 and AKR1C3 co-expression

HSD11B1 and AKR1C3 transfections were performed as described above. Transfected cells were re-plated at a 1:1 ratio of HSD11B1:AKR1C3 (2×10^5^ cells/mL, 300 µL) in 48-well Corning® CELLBIND® surface plates. After 24 hours, media was replaced with serum free media containing 100 nM cortisone in combination with either 10 nM or 100 nM 11KA4 in the absence and presence of 10 µM CBX. Aliquots (200 µL) were collected over a 24 hour period for analysis by UHPLC-MS/MS.

### Promoter reporter assays

To determine the effect of the HSD11B1 inhibitor CBX on AR and GR transactivation, HSD11B1 and AKR1C3 transfections were performed 24 hours before co-transfections of HEK293 cells with either a AR expression plasmid (0.09 µg) and the pTAT-GRE-Elb-luc luciferase promoter reporter construct (0.9 µg) or a GR expression plasmid (0.09 µg) and the pTAT-GRE-Elb-luc luciferase promoter reporter construct (0.9 µg). HSD11B1 and AKR1C3 transfected cells were re-plated at a 1:1 ratio (4×10^5^ cells/mL, 1 mL) in 12-well Corning® CELLBIND® surface plates. Following 24 hours, media was replaced with serum free media containing 100 nM cortisone and 10 nM 11KA4 in the absence and presence of 10 µM CBX. At the same time the AR-GRE transfected cells and GR-GRE transfected cells were re-plated into 48-well Corning® CELLBIND® plates (3×10^5^ cells/mL, 300 µL). Following a 24 hour incubation with steroid, aliquots (200 µL) from the HSD11B1:AKR1C3 expressing cells were collected for analysis of steroid metabolism by UHPLC-MS/MS. A further 300 µL conditioned media was transferred to each of the AR-GRE and GR-GRE transfected cells. These cells were incubated for a further 24 hours after which cell lysates were harvested in passive lysis buffer. Reporter activity was then measured by making use of the Luciferase Assay System (Promega). The luciferase values were normalised to the protein concentration of each individual sample determined using a Thermo Scientific™ Pierce™ BCA Protein Assay kit.

### Determining the effect of ADZ4017 on 11KA4 conversion in human adipose tissue

Paired subcutaneous and omental adipose tissue was collected from eight female participants (age range 32-59 years, BMI range 42-57 kg/m^2^) undergoing elective abdominal non-cancer surgery (Black Country Research Ethics Committee references 14/WM/0011 and 19/WM/0183). Written, informed consent was obtained prior to surgery and sample collection. Adipose tissue samples were transported to the laboratory at room temperature in DMEM/F12 supplemented with 33 µM biotin, 17 µM panthotenic acid and 1% penicillin-streptomycin. Connective tissue and vessels were removed and the adipose tissue washed in PBS. Tissue samples of 100-300 mg were cut and weighed. Each sample was cut into four smaller pieces for incubations with 100 nM cortisone or 100 nM 11KA4 in the presence and absence of 100 nM HSD11B1 inhibitor AZD4017 in DMEM/F12-Ham (supplemented as described above). Final solvent concentrations were adjusted to 0.00302% MeOH and 0.1% DMSO in all incubations. Tissue incubations were constantly rotated in a hybrid oven at 37 °C for 72 hours. Medium samples were centrifuged for 10 minutes at 16,000g and 4 °C and the supernatant was used for analysis of steroid metabolism by UHPLC-MS/MS. The tissue was washed in PBS, snap frozen in liquid nitrogen and stored at −80 °C for RNA extraction.

### RNA isolation and gene expression analysis from HEK293 cells

Total RNA was isolated from cells using Tri-Reagent® (Sigma) and cDNA synthesis carried out using the GoScript™ Reverse Transcription System kit (Promega). Quantitative PCR was performed using a Roche LightCycler® 96 rapid thermal cycler instrument and the KAPA SYBR® FAST qPCR Master Mix for LightCycler®. Primer sequences as follows, AKR1C3 (forward) 5’-AGCCAGGTGAGGAACTTTC-3’ (53), AKR1C3 (reverse) 5’- ATCACTGTTAAAATAGTGGAG-3’ (53), HSD11B1 (forward) 5’- CTGCCTGCTTAGGAGGTTGT-3’, HSD11B1 (reverse) 5’- CCTTGGAGCATCTCTGGTCTG-3’, the reference gene ubiquitin conjugating enzyme (UBC) (forward) 5’- CCGGGATTTGGGTCGCAG-3’, UBC (reverse) 5’- TCACGAAGATCTGCATTGTCAAG-3’. Primer efficiencies were between 90% and 110% for all primer sets. Relative gene expression was calculated using a modified form of the model described by Pfaffl (54). Inter-experimental variations were adjusted using a mean-centering (55).

### RNA isolation and gene expression analysis from adipose tissue

Adipose tissue was homogenised in Tri-Reagent® and total RNA was isolated after phenol-chloroform extraction using the Qiagen RNeasy Mini Kit using an established protocol (56). Reverse transcription was performed using Applied Biosystems™ TaqMan™ Reverse Transcription Reagents according to the manufacturer’s protocol. qPCR was performed on an ABI 7900HT sequence detection system (Perkin Elmer, Applied Biosystems) using TaqMan™ Gene Expression Assays (FAM-labelled) and the SensiFAST^TM^ Probe Hi-ROX kit (Bioline). The following TaqMan™ Gene Expression Assays were used: 18S Hs99999901_s1, AKR1C3 Hs00366267_m1, HSD11B1 Hs01547870_m1.

### Determining the effect of ADZ4017 glucocorticoid and androgen concentrations in vivo

A total of 28 participants (22 male and 6 female), all with type 2 diabetes, were recruited for a randomized, double-blind, parallel-group, placebo-controlled trial. Full ethical approval was acquired from North West Greater Manchester Central Research Ethics Committee 17/NW/0283 prior to the initiation of recruitment for the study. Informed consent was obtained from all participants after the nature and possible consequences of the study had been explained. Participants received either oral AZD4017 (400 mg) or a matched oral placebo twice daily for 35 days. Blood samples and urine were collected from individuals before first treatment and after completion of the trial and were analysed by UHPLC-MS/MS and GC-MS, respectively. Refer to Ajjan *et al.* (40) for further details on study design and participants.

### Steroid extraction

Steroids were extracted from HEK293 tissue culture media using Tert-butyl methyl ether (MTBE) (57). In brief, following the addition of an internal standard mix (100 µL containing 2 ng D3-T and 5 ng D7-A4, D7-11OHA4, D3-11KT and D4-F in acetonitrile) and MTBE (3:1 ratio of MTBE:media), samples were vortexed at 1 500 rpm for 10 minutes, after which they were incubated at −80 °C for 1 hour, allowing for the aqueous layer to freeze. The resulting organic phase was subsequently transferred to a clean test tube and evaporated under a stream of nitrogen gas at 45 °C. Extracted steroids were then reconstituted by the addition of 50% (v/v) methanol and stored at −20 °C until analysis by UHPLC-MS/MS. Serum steroids and steroids in medium adipose tissue incubations were extracted using a similar method. Samples (200 μL for serum, 500 μL for adipose tissue conditioned medium) were mixed with 10 μL internal standards in 50% (v/v) methanol in water (D4-F, D8-E, D6-DHEA, D7-A4, D3-T, D7-11OHA4, D10-11KA4, D3-11KT; 5 ng each for serum, 20 ng each for adipose tissue). For serum samples 50 μL acetonitrile were added for protein precipitation prior to extraction and the samples vortexed. MTBE was added to each sample (1 mL for serum; 3 mL for medium from adipose tissue incubations) and the samples were vortexed at 1000 rpm for 10 min. Following incubation at room temperature for 30 minutes the organic phase was transferred, dried under a stream of nitrogen gas at 45 °C and reconstituted with 50% (v/v) methanol in water prior to analysis by UHPLC-MS/MS. For the measurement of serum DHEAS, 200 ng D2-DHEAS in 20 uL of 50% methanol was added to 20 μL of serum followed by 20 μL of 1 mM ZnSO4 and 100 μL acetonitrile. The samples were centrifuged and 100 μL of the supernatant was dried and reconstituted in 50% (v/v) methanol in water prior to UHPLC-MS/MS analysis.

### Quantification of steroids by ultra-high performance liquid chromatography tandem mass spectrometry (UHPLC-MS/MS)

Steroids from assays performed in HEK293 cells were separated and quantified using a UPLC high strength silica (HSS) T3 column (2.1 mm X 50 mm, 1.8 µm particle size) coupled to an ACQUITY UPLC and Xevo TQ-S triple quadrupole mass spectrometer (Waters Corporation, Milford, USA) as previously reported (57). Steroids extracted from serum or adipose tissue incubations were separated and quantified using Phenomenex Luna Omega polar C18 column (2.1 × 50 mm, 1.6 µm particle size; Phenomenex, Macclesfield, UK) coupled to an ACQUITY UPLC and a Waters Xevo TQ-XS triple quadrupole mass spectrometer as previously described (58). Serum DHEAS was quantified using a HSS T3 column (2.1 × 50 mm, 1.8 µm particle size; Waters) coupled to an ACQUITY UPLC and a Waters Xevo TQ-XS triple quadrupole mass spectrometer as previously described (59, 60).

### Quantification of urinary steroids by GC-MS

Measurement of urinary steroid metabolite excretion was carried out by a long-established gas chromatography-mass spectrometry (GC-MS) method (61). In brief, free and conjugated steroids were extracted from 1 mL urine by solid-phase extraction. Steroid conjugates were enzymatically hydrolyzed, reextracted, and chemically derivatized to form methyloxime trimethyl silyl ethers. An Agilent 5975 instrument operating in selected-ion-monitoring (SIM) mode achieved sensitive and specific detection and quantification of selected steroid metabolites.

### Statistical analysis

We performed unpaired t-tests to compare the effect of HSD11B1 inhibition on 11-oxygenated androgen production in HEK293 cells expressing AKR1C3 and HSD11B1 and on the transactivation of the androgen receptor. Paired t-tests were performed to assess the effect of HSD11B1 inhibition in adipose tissue, while Wilcoxon signed-rank tests were used to assess the effect in serum and urine. Correlations between serum steroid concentrations and clinical outcomes were investigated using a Spearman’s Rho test.

## Results

### HSD11B1 efficiently catalyses the 11β-reduction of 11-oxygenated androgens

Non-steroidogenic HEK293 cells were transiently transfected to express HSD11B1 and activity assayed. Rate equations were fitted to the progress curves for three independent experiments as shown in **Supplementary** Figure 1. A low but detectable reverse reaction rate was observed for the cortisone to cortisol and the 11KA4 to 11OHA4 reactions. Reversible kinetics were therefore used for these reactions. Our results show that the apparent Km (Km, *app*) of HSD11B1 for cortisone and 11KA4 are similar (0.9 µM and 1 µM), while the Km, *app* for 11KT (0.21 µM) is >4-fold lower, indicating a higher affinity towards 11KT (**Table 1**). However, the apparent Vmax (Vmax, app) for 11KT (0.05 μM/h) was >2-fold lower than that for cortisone (0.12 μM/h) and 7-fold lower than that for 11KA4 (0.35 μM/h). Considering the Vmax/Km values, an estimate of enzyme efficiency, both 11KA4 and 11KT are more efficiently converted to their respective products than the traditionally recognised HSD11B1 substrate cortisone.

**Table 1:** Kinetic constants (K_m, *app*_; _Vmax, *app*;_ V_max_/K_m_) determined for HSD11B1.

| Reaction | $K_{m, app}$ (substrate)<br>($\mu M$) | $K_{m, app}$ (product)<br>($\mu M$) | $V_{max, app}$ (forward)<br>($\mu M/h$)<sup>*</sup> | $V_{max, app}$ (reverse)<br>1/h<sup>a</sup> or ($\mu M/h$)<sup>b</sup> | $V_{max}/K_m$<sup>#</sup> |
| --- | --- | --- | --- | --- | --- |
| Cortisone -> Cortisol | 0.9 | - | 0.12 | 0.01<sup>a</sup> | 0.13 |
| 11KA4 -> 11OHA4 | 1 | 2.46 | 0.35 | 0.03<sup>b</sup> | 0.35 |
| 11KT -> 11OHT | 0.21 | - | 0.05 | - | 0.24 |

<sup>\*</sup> $V_{max, app}$ values are dependent on transfection efficiency and differ between experiments. We included the conversion of 100 nM 11KA4 to 11OHA4 in all experiments and used the initial rate to normalise the transfection efficiency, which was set to 1 for the values reported above.
<sup>#</sup> $V_{max}/K_m$ values are reported for the substrate and forward reaction.

### Co-expression of HSD11B1 with AKR1C3 modulates the activation of 11-oxygenated androgens

AKR1C3 is the key enzyme catalysing the peripheral conversion of androgen precursors into active androgens (1>). We have previously shown that AKR1C3 catalyses the conversion of 11KA4 to the potent androgen 11KT 8-fold more efficiently than the conversion of androstenedione to testosterone (16). To determine how the co-expression of HSD11B1 with AKR1C3 would affect 11KT production, we mixed transfected HEK293 cells expressing either AKR1C3 or HSD11B1 to obtain increasing ratios of HSD11B1:AKR1C3, while keeping the number of cells expressing AKR1C3 constant. The total cell count was also kept constant by including cells transfected with an empty plasmid as required. The relative expression of HSD11B1:AKR1C3 was confirmed by qPCR (**Figure 2A**). Increased ratios of HSD11B1:AKR1C3 resulted in increased concentrations of cortisol being produced upon addition of 100 nM cortisone (**Figure 2B**), thereby confirming the increased HSD11B1 expression. Similarly, increasing the ratio of HSD11B1:AKR1C3 resulted in the concomitantly increased production of 11OHA4 from 11KA4, with 11OHA4 becoming the most abundant product when the HSD11B1:AKR1C3 ratio was increased beyond 0.1:1 (**Figure 2C**). Biosynthesis of 11KT was highest when AKR1C3 was expressed in the absence of HSD11B1, with increasing HSD11B1:AKR1C3 ratios resulting in a rapid decrease in the concentrations of 11KT. 11OHT initially increased as the ratio of HSD11B1:AKR1C3 increased, peaking at a ratio of 0.25:1. Further increases in the HSD11B1:AKR1C3 ratio resulted in decreased concentrations of 11OHT being produced.

**Figure 2:**
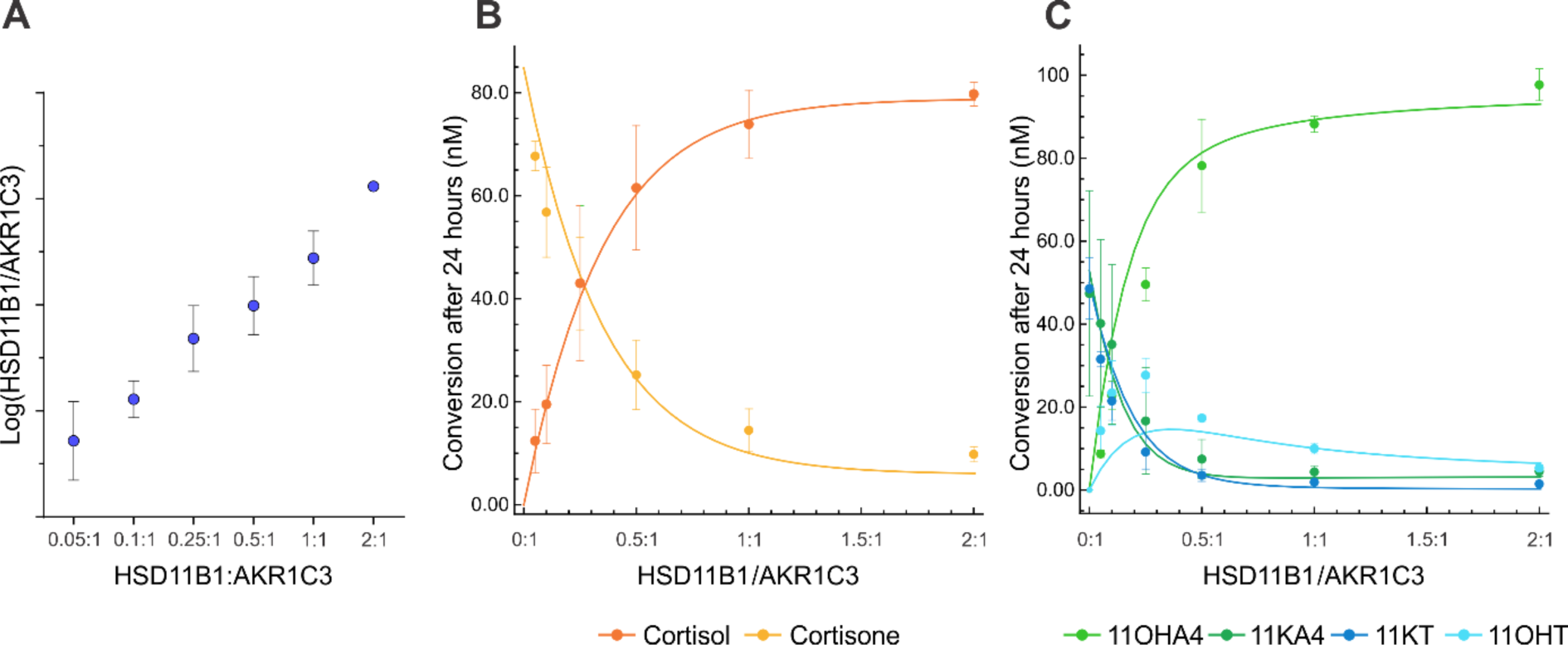
Increased HSD11B1:AKR1C3 ratios modulate the biosynthesis of 11KT. **(A)** qPCR data confirming the increase in HSD11B1 expression relative to that of AKR1C3 at the different HSD11B1:AKR1C3 ratios. **(B)** The conversion of cortisone (0.1 μM) to cortisol by increasing ratios of HSD11B1:AKR1C3 after 24 hours **(C)**. The conversion of 11KA4 (0.1 μM) to 11OHA4, 11KT and 11OHT by increasing ratios of HSD11B1:AKR1C3 after 24 hours. For **B** and **C** the experimental data from the ratio experiments are shown as points with error bars, while the computational models’ predictions are shown with the solid connecting lines. All results are shown as means ± SEM of three independent experiments.

Using the parameterised rate equations for HSD11B1 and those previously determined by us for AKR1C3 (1>), a computational model consisting of a set of ordinary differential equations was constructed to simulate steroid metabolism in cells with both enzymes expressed:

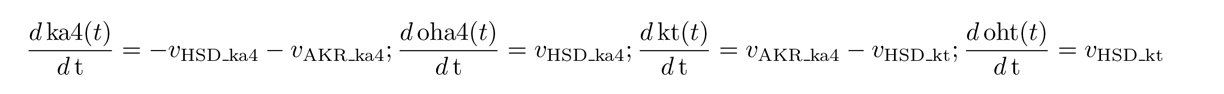

The rate equations for HSD11B1 and AKR1C3 are given in the Supplementary material. The model was able to predict the experimentally determined effect of increasing HSD11B1:AKR1C3 ratios on the metabolism of cortisone and 11KA4, respectively (**Figure 2**). Note that the kinetic parameters in the mathematical model were not fitted to the experimental data for the different HSD11B1:AKR1C3 ratios, but were determined in the progress curves for the individual incubations (**Supplementary** Figure 1). Taken together these data clearly show that HSD11B1 modulates the biosynthesis of 11KT by AKR1C3, and that the model can predict the effect of varying HSD11B1:AKR1C3 ratios.

### In vitro inhibition of HSD11B1 results in the accumulation of 11-ketotestosterone

Next, we set out to determine the effect of HSD11B1 inhibition on 11KT biosynthesis. We mixed HEK293 cells transiently transfected with either AKR1C3 or HSD11B1 at a ratio of 1:1 and simultaneously treated the cells with cortisone and 11KA4 in the absence and presence of the HSD11B1 inhibitor, carbenoxolone (CBX). We used either equimolar concentrations of cortisone and 11KA4 (100 nM each) or 10-fold excess of cortisone (100 nM) over 11KA4 (10 nM) to more closely mimic circulating concentrations (>). Our results show that CBX reduced the production of cortisol from cortisone from 56.5 nM to 3.2 nM after 24 hours, demonstrating effective HSD11B1 inhibition (**Figure 3A** **& B**). In the absence of CBX the majority of 11KA4 (69%) was converted to 11OHA4 by HSD11B1, with only 3.8 nM of 11KT produced from 100 nM 11KA4 by AKR1C3 (**Figure 3C**). However, inhibition of HSD11B1 resulted in a 7-fold increase in 11KT production after 24 hours (**Figure 3D**). Similarly, in cells treated with 100 nM cortisone and 10 nM 11KA4, inhibition of HSD11B1 by CBX resulted in a 12-fold increase in 11KT production (0.15 nM without CBX to 1.8 nM with CBX) (**Figure 3E** **& F**). 11KT was the by far most abundant 11-oxygenated androgen following inhibition of HSD11B1. Inhibition of both the conversions from 11KA4 to 11OHA4 and 11KT to 11OHT, respectively, allowed AKR1C3 to catalyse the conversion of 11KA4 to 11KT without the need to compete for substrate and prevented the inactivation of 11KT to the less potent 11OHT. The computational model was able to independently predict the outcome of HSD11B1 inhibition under the experimental conditions (**Figure 3**).

**Figure 3:**
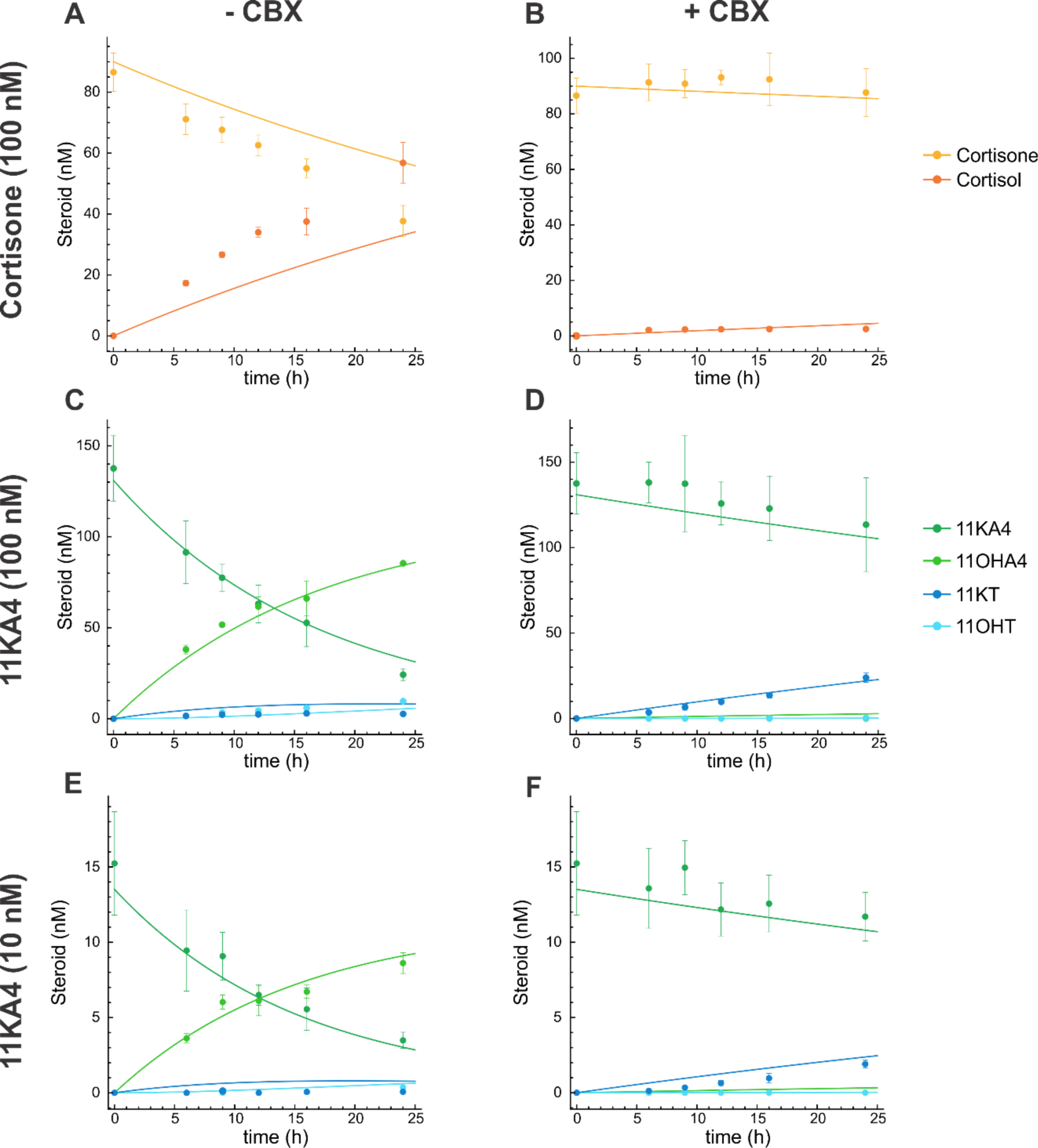
Inhibition of HSD11B1 by carbenoxolone (CBX) inhibits the biosynthesis of cortisol, but allows for increased AKR1C3-mediated production of 11KT when HSD11B1 and AKR1C3 are co-expressed. HEK293 cells expressing HSD11B1 and AKR1C3 were combined at a ratio of 1:1 and treated with either 100 nM cortisone **(A & B)** and 100 nM 11KA4 **(C & D)** with and without CBX (10 µM) or 100 nM cortisone and 10 nM 11KA4 **(E & F)** with or without CBX (10 µM). The experimental data from the time course experiment is shown as points with error bars, while the computational models’ predictions are shown with the solid lines. All results are shown as means ± SD of a single biological experiment performed in triplicate. This enabled the model to predict the outcomes from one experiment with the same transfection efficiency.

### In silico analysis of the combined effect of HSD11B1:AKR1C3 co-expression and HSD11B1 inhibition on 11KT biosynthesis

After having shown that the computational model was able to individually predict both the effect of increasing HSD11B1:AKR1C3 ratios (**Figure 2**) and the effect of HSD11B1 inhibition (**Figure 3**), we used the model to simultaneously predict the effect of differing HSD11B1 and AKR1C3 expression and differing degrees of HSD11B1 inhibition on the conversion of 10 nM 11KA4 to 11OHA4, 11KT and 11OHT after 24 hours (**Figure 4**). These results confirmed that HSD11B1 strongly modulates the biosynthesis of 11KT by competing for the precursor, 11KA4, and by converting the 11KT that is formed to the less potent androgen 11OHT. The inhibition of HSD11B1 reduced these effects under all HSD11B1:AKR1C3 ratios tested leading to increased production of 11KT. Notably, lower ratios of HSD11B1:AKR1C3 were more sensitive to HSD11B1 inhibition, more quickly leading to increased 11KT concentrations as the level of inhibition was increased. Higher absolute concentrations of 11KT were also produced by lower ratios of HSD11B1:AKR1C3 (AKR1C3 was kept constant) when HSD11B1 was inhibited up to the maximum simulated I/Ki value of ten. Near complete inhibition of HSD11B1 (I/Ki = 100), however, negated the effect of the HSD11B1:AKR1C3 ratio with 11KT production dependent only on AKR1C3 (**Supplementary** Figure 2). The effect of HSD11B1 inhibition is therefore dependent on the degree of inhibition, the ratio of HSD11B1:AKR1C3 and the absolute expression level of HSD11B1 and AKR1C3.

**Figure 4:**
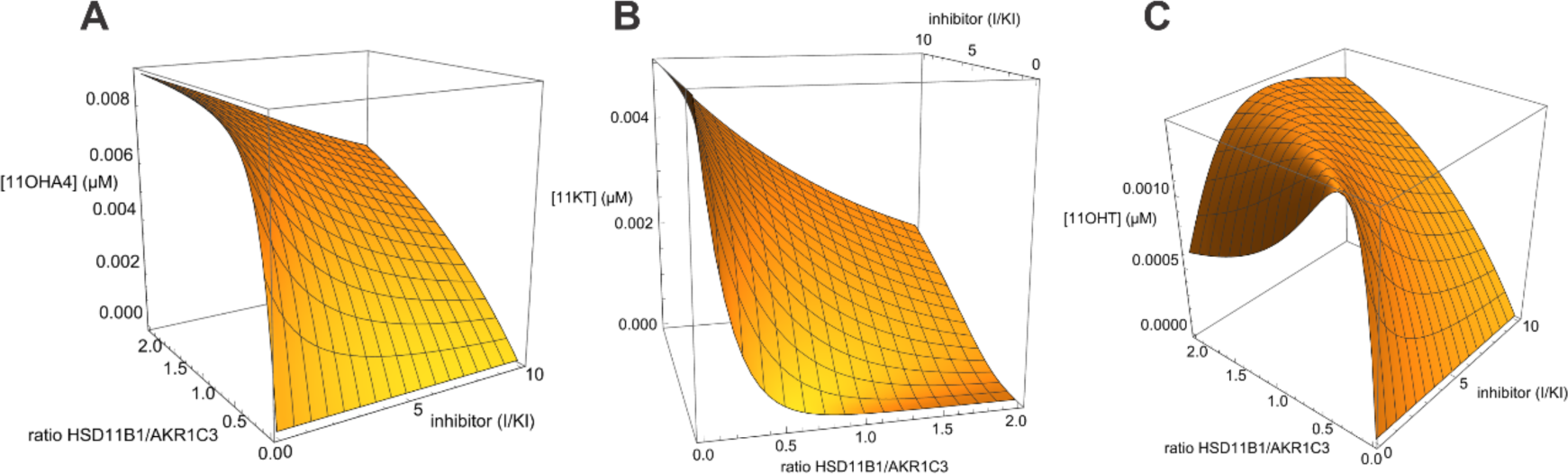
Computational model of the effect of varying HSD11B1:AKR1C3 ratios and different levels of HSD11B1 inhibition on the resulting concentrations of 11OHA4 (A), 11KT (B) and 11OHT (C). The model varies that ratio of HSD11B1:AKR1C3 as in Figure 2 and estimates the effect of different levels of HSD11B1 inhibition on the conversion of 10 nM 11KA4 after 24 hours as in Figures 2 and 3.

To investigate the potential biological impact of HSD11B1 inhibition, we combined HEK293 cells expressing AKR1C3 or HSD11B1 at a 1:1 ratio and incubated the cells with 100 nM cortisone and 10 nM 11KA4 in the absence and presence of CBX for 24 hours. The conditioned media were then transferred to HEK293 cells transiently transfected with either a plasmid expressing the glucocorticoid receptor (GR) or the androgen receptor (AR), and a luciferase reporter construct. The GR-and AR-mediated induction of luciferase was subsequently measured after an additional 24 hour incubation. As expected, HSD11B1 mediated the conversion of cortisone to cortisol, with the resulting cortisol increasing the transactivation of the GR (94-fold induction). Inhibition of HSD11B1 with CBX reduced cortisol production 32-fold (**Figure 5A**) and resulted in a 34-fold decrease in GR transactivation (**Figure 5B**). Conversely, inhibition of HSD11B1 resulted in a 7-fold increase in transactivation of the AR due to significantly increased 11KT biosynthesis (**Figure 5C** **& D**). Basal levels of AR transactivation in the absence of CBX and 11KT were due to the presence of low concentrations of 11OHT (0.81 nM; **Supplementary** Figure 3), which is a partial agonist of the AR (11, 62). It should be noted that we observed that CBX weakly antagonises the AR (**Supplementary** Figure 4), hence, the induction of the AR was likely underestimated. Nonetheless, taken together, these data demonstrate that while inhibition of HSD11B1 results in the desired decrease in GR transactivation, this is accompanied by a shift towards AR transactivation.

**Figure 5:**
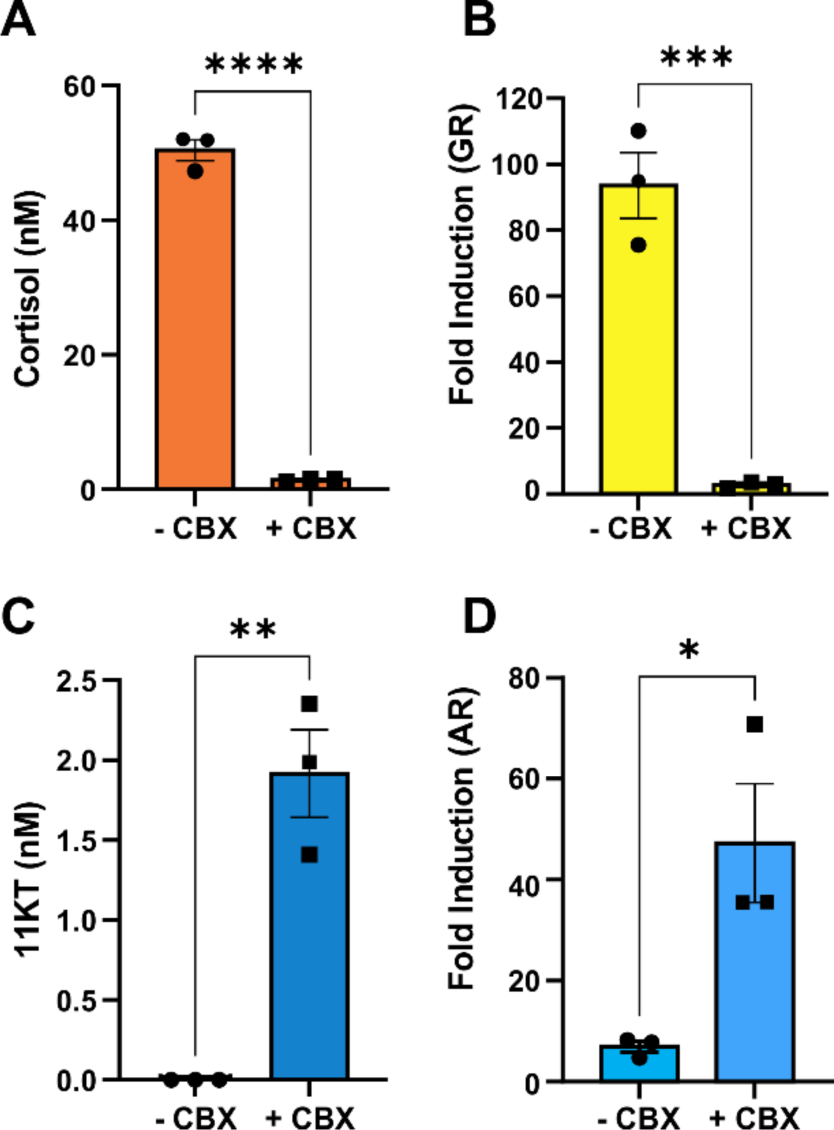
Inhibition of HSD11B1 by carbenoxolone (CBX) results in decreased cortisol production (A) and an accompanying decrease in transactivation of the glucocorticoid receptor (B), but increased production of 11KT (C) and transactivation of the androgen receptor (D). HEK293 cells expressing HSD11B1 and AKR1C3 were combined at a ratio of 1:1 and treated with 100 nM cortisone and 10 nM 11KA4 with or without CBX (10 µM). After 24 hours the conditioned media was transferred to HEK293 cells transiently transfected with either a plasmid expressing the glucocorticoid receptor (GR) and a luciferase reporter construct or a plasmid expressing the androgen receptor (AR) and the same luciferase reporter construct. Luciferase was assayed after an additional 24 hour incubation and is shown as the fold-induction relative to a vehicle control. All results are shown as means ± SEM of three biological experiments. P values were calculated using unpaired t-tests (*P <0.05; **P<0.01; ***P<0.001; ****P <0.0001).

### Selective HSD11B1 inhibition increases 11KT production in female human adipose tissue ex vivo

We next set out to determine the effect of HSD11B1 inhibition on 11KT biosynthesis in paired subcutaneous and omental adipose tissue samples from women undergoing elective bariatric surgery. For these experiments we employed the selective HSD11B1 inhibitor AZD4017 (63). The conversion of cortisone to cortisol was reduced to 3% of the uninhibited conversion rate in subcutaneous adipose tissue treated with AZD4017, thereby confirming effective HSD11B1 inhibition (**Figure 6A**). Following incubation of subcutaneous adipose tissue with 11KA4 (100 nM), the most abundant product formed in the absence of AZD4017 was 11OHT (60.1% of total products formed), followed by 11OHA4 (23.4%) and 11KT (16.5%) (**Figure 6F**). The biosynthesis of both 11KT and 11OHT is dependent on AKR1C3 activity and the observed sum of 11KT and 11OHT (76.6% of products formed) suggested substantial AKR1C3 expression. This was confirmed by qPCR analysis, which showed higher AKR1C3 expression relative to that of HSD11B1 (**Figure 6C**). However, HSD11B1 converted 78.5% of the 11KT formed to the less active 11OHT. Inhibition of HSD11B1 by AZD4017, resulted in a 5-fold increase in 11KT, with 11KT becoming the most abundant product (85.6%) (**Figure 6G**). This was accompanied by significant decreases in the production of both 11OHA4 and 11OHT (21-fold and 4.5-fold, respectively).

**Figure 6:**
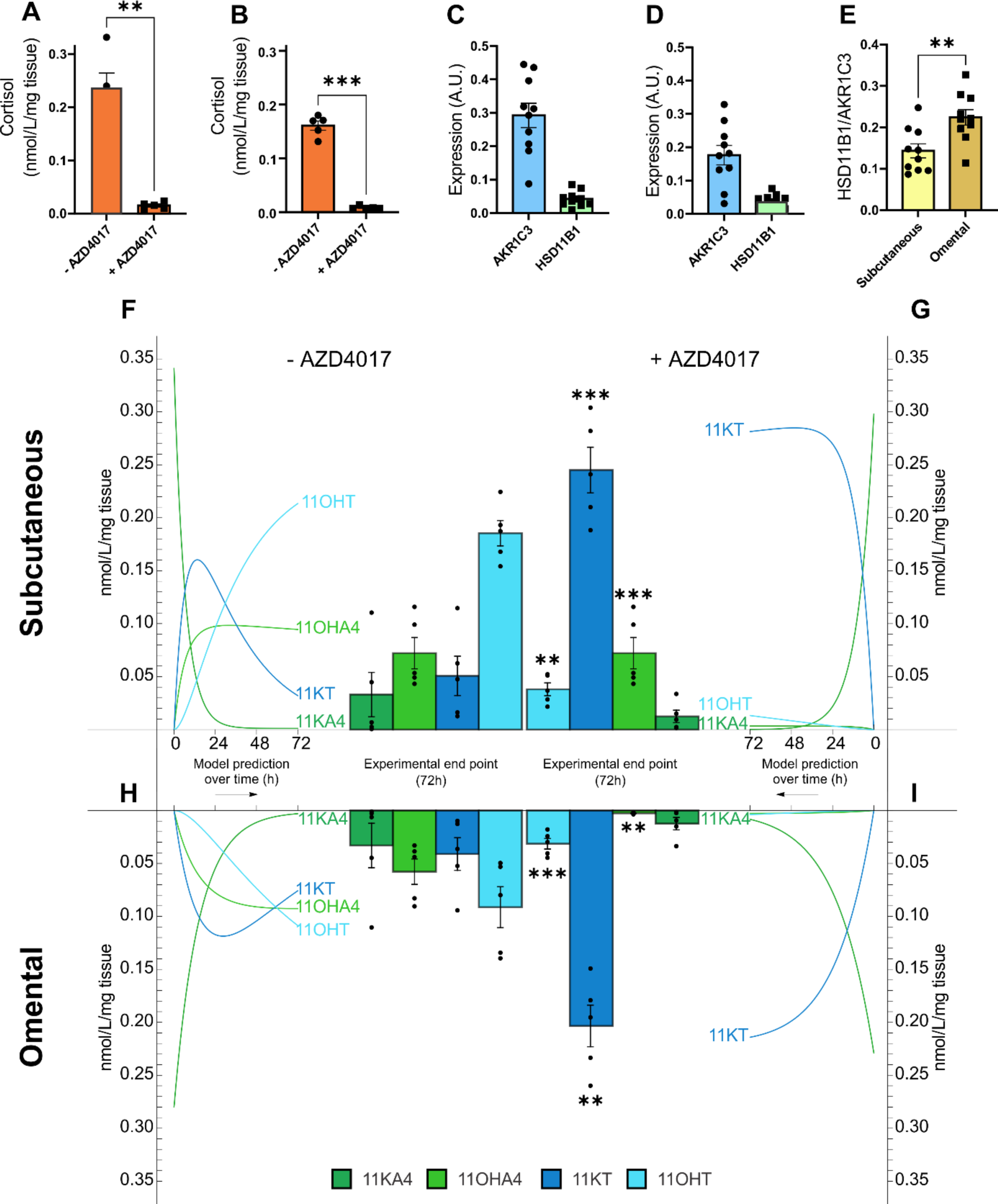
Inhibition of HSD11B1 by AZD4017 in human subcutaneous (A, F, G) and omental (B, H, I) adipose tissue results in the increased production of 11KT. Adipose tissue was incubated *ex vivo* with 100 nM cortisone (**A, B**) or 11KA4 (**F-I**) with or without AZD4017 (100 nM) for 72 hours. The mathematical model’s prediction of the conversion over time is shown next to the experimentally determined end point data after the 72 hour incubation. Expression of AKR1C3 and HSD11B1 in subcutaneous (**C**) and omental (**D**) adipose tissue was measured by qPCR. The expression ratio of HSD11B1:AKR1C3 were determined for both adipose tissue subtypes (**E**). P values were calculated using Wilcoxon signed-rank tests (*P <0.05; **P<0.01; ***P<0.001).

Marginally lower conversion of cortisone to cortisol was observed in omental adipose tissue as compared to subcutaneous tissue, with AZD4017 again effectively inhibiting HSD11B1 activity (**Figure 6B**). Treatment with AZD4017 again resulted in a significant decrease in 11OHA4 and 11OHT concentrations, with an accompanying 4-fold increase in 11KT production (**Figure 6H** **& I**).

The sum of 11KT and 11OHT (66.0%) was lower in the omental tissue than in subcutaneous tissue (76.6%) illustrating lower AKR1C3 expression as confirmed by qPCR (**Figure 6C** **& D**). The ratio of HSD11B1 over AKR1C3 expression was lower in subcutaneous than omental adipose tissue (**Figure 6E**), favouring androgen activation in subcutaneous tissue vs glucocorticoid activation in omental tissue. The metabolism of 11KA4 and the effect of AZD4017 inhibition in human adipose tissue was simulated with the mathematical model that was constructed using conversion data obtained from transfected HEK293 cells. The expression levels of the HSD11B1 and AKR1C3 in the adipose tissue relative to that in transfected HEK293 cells therefore needed to be estimated. This was achieved by fitting the expression levels of HSD11B1 to the cortisone to cortisol conversion data (**Figure 6A** **& B**) and then using an AKR1C3 activity based on the AKR1C3/HSD11B1 expression ratio as determined by qPCR (**Figure 6C** **& D**). This allowed the model to predict the conversion of 11KA4 to the respective products over time. These simulation revealed the initial accumulation of 11KT, prior to the time dependent conversion of this product to 11OHT by HSD11B1 (**Figure 6F** **&H**). Inhibition of HSD11B1 eliminated the latter conversion yielding only 11KT and illustrated that the amount of 11KT formed over time was dependent on the absolute expression levels of AKR1C3.

Taken together, these data indicate that inhibition of HSD11B1 in adipose tissue leads to a shift in the metabolism of 11-oxygenated androgens resulting in increased 11KT biosynthesis.

### In vivo treatment with a selective HSD11B1 inhibitor increases circulating 11KT concentrations

After having shown that inhibition of HSD11B1 in adipose tissue shifts the metabolism of 11-oxygenated androgens towards 11KT biosynthesis, we next set out to determine if inhibition of HSD11B1 would alter circulating concentrations of 11KT. We therefore measured a panel of 11-oxygenated androgens as well as cortisol and cortisone in the serum of individuals with type 2 diabetes who had received treatment with the selective HSD11B1 inhibitor AZD4017 during a randomized, double-blind, parallel-group, placebo-controlled phase 2b pilot trial (40). Paired (day 0 and day 35) samples were available for 18 participants (13 men and 5 women; median BMI 32.2 kg/m^2^ range 25.0-47.5 kg/m^2^; median age 64.5, range 28-84 years), with 11 randomised to HSD11B1 inhibitor treatment (7 men and 4 women) and 7 (6 men and 1 woman) to placebo.

Our measurements confirmed that AZD4017 significantly reduced the serum cortisol/cortisone ratio, indicating effective HSD11B1 inhibition as previously reported for selected urinary glucocorticoid metabolites by LC-MS/MS (40), and further confirmed here by GC-MS **(Supplementary** Figure 5). However, concomitantly, circulating 11KT concentrations significantly increased 2.5-fold (from 1.7 to 4.3 nM) following treatment with 400 mg AZD4017 twice daily for 35 days. This was accompanied by a significant, 2-fold decrease in 11OHT concentrations and a 4-fold decrease in the 11OHT/11KT ratio (**Figure 7**). While the circulating concentrations of the 11-oxygenated androgen precursors 11OHA4 and 11KA4 did not change, a small, but significant decrease was also observed for the 11OHA4/11KA4 ratio. These changes were also reflected by a significant increase in the urinary metabolite 11-oxoandrosterone, which is derived from 11KA4 and 11KT (64), and a 6-fold decrease in the 11β-hydroxyandrosterone/11-oxoandrosterone ratio (**Figure 7**). No changes in the serum concentrations of 11-oxygenated androgens were observed in the placebo-treated group (**Supplementary** Figure 6).

**Figure 7:**
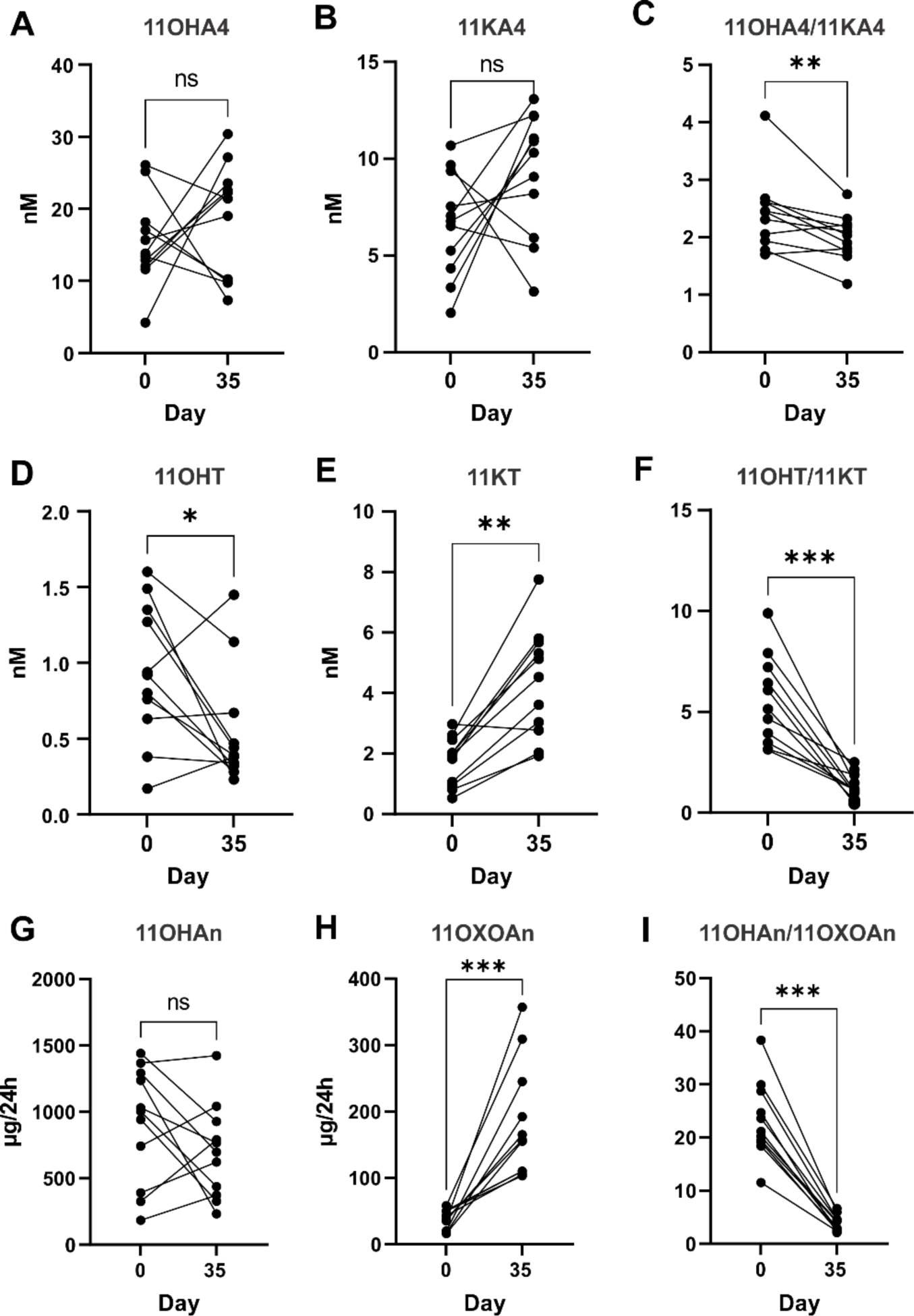
Inhibition of HSD11B1 by oral administration of AZD4017 (400 mg twice daily) for 35 days (n=11) changes the serum (A-F) and urinary (G-I) profile of 11-oxygenated androgens and their metabolites, with significant increases in the circulating concentration of the potent androgen 11KT. P values were calculated using Wilcoxon signed-rank tests (*P <0.05; **P<0.01; ***P<0.001). Serum concentrations of classic androgens are shown in **Supplementary** Figure 7.

When considering correlations between the changes in circulating steroid concentrations, it is interesting to note that the cortisol/cortisone ratio is highly correlated with the change in both the ratios of 11OHA4/11KA4 (R = 0.893; P < 0.001) and 11OHT/11KT (R = 0.686; P < 0.01), but not with the change in circulating 11KT (R = −0.375; P = 0.126) (**Supplementary Table 1**).

## Discussion

The relatively recent identification of 11-oxygenated androgens as important contributors to the human androgen pool has prompted a re-evaluation of existing paradigms. One such paradigm is the idea that the two HSD11B isoforms are predominantly involved in the peripheral metabolism of glucocorticoids. We and others have previously shown that HSD11B1 and HSD11B2 are essential for the interconversion of the 11β-hydroxyl and 11-keto forms of the 11-oxygenated androgens (11, 20). We have also previously shown that 11KA4, but not 11OHA4, is a substrate for the key androgen-activating enzyme AKR1C3 (16, 17). Therefore, HSD11B2 plays a vital role in catalysing the conversion of adrenal-derived 11OHA4 to 11KA4, which can then be converted to the potent androgen 11KT by AKR1C3 (13). One of the main sites of AKR1C3 expression and peripheral androgen activation is adipose tissue (14, 15) which also abundantly expresses HSD11B1 (19, 22, 65). Paulukinas et. al. have previously observed that co-expression of HSD11B1 with AKR1C3 in differentiated SGBS adipocyte cells may prevent the accumulation of 11KT, thus suggesting that HSD11B1 may protect from androgen excess (17). Here, using a combination of *in vitro*, *in silico* and *ex vivo* studies, we demonstrate that co-expression of HSD11B1 with AKR1C3 modulates peripheral biosynthesis of 11KT, first, by competing with AKR1C3 for the substrate 11KA4, and, second, by converting much of the resulting 11KT to the less potent androgen 11OHT. We demonstrated that inhibition of HSD11B1 results in significantly increased 11KT biosynthesis as shown by us *in vitro*, *in silico*, *ex vivo* and in a phase 2a study comparing the effects of 35 days of treatment with the selective HSD11B1 inhibitor AZD4017 to placebo treatment. Our data show that HSD11B1 inhibition does not only alter local, adipose tissue-specific 11KT concentrations, but also systemic levels, thus emphasising that peripheral tissues expressing AKR1C3 are essential for 11KT biosynthesis (13). Collectively, our findings show that under normal circumstances HSD11B1 plays an important novel role in preventing 11KT accumulation, in addition to its well-known role in local glucocorticoid activation.

Our findings are of immediate clinical relevance, particularly in women, where circulating concentrations of 11KT are similar to testosterone concentrations (3, 6, 66, 67). Therefore, increasing 11KT due to HSD11B1 inhibition markedly increases the total circulating pool of active androgens in women. This increase in androgen load is significant given that 11KT has shown functional activity comparable to testosterone (5, 62, 68) and increased circulating androgens have been linked to a significantly greater risk of type 2 diabetes in women (43). Moreover, there is growing evidence that androgens drive the risk of common metabolic disorders, including type 2 diabetes, hypertension, cardiovascular disease and MASLD (15, 43, 69–72, 44, 46–52), in women with PCOS where androgen excess is a hallmark (72). Notably, O’Reilly et. al. have previously shown that local AKR1C3-mediated androgen activation in adipose tissue drives lipotoxicity in women with PCOS (15). They propose a vicious cycle in which increased androgen generation in adipose tissue results in lipid accumulation, increased fat mass and as a result systemic insulin resistance. This cycle is completed by the insulin-induced stimulation of AKR1C3 expression in adipose, leading to increased androgen activation. Given that 11KA4 is the preferred substrate for AKR1C3, increased insulin-induced AKR1C3 expression would therefore lead to significantly more 11KT being produced, which would be further increased by HSD11B1 inhibition (16, 17, 73).

Our finding of increased 11KT biosynthesis to clinically highly relevant concentrations occurring after HSD11B1 inhibition therefore needs to be taken into account when considering HSD11B1 inhibition as treatment for type 2 diabetes or metabolic syndrome in women, as the beneficial effects of reduced local glucocorticoid activation could be offset by opposing metabolic effects due to increased 11KT concentrations and AR signalling. This is highly relevant for the recently proposed use of HSD11B1 inhibitors in ameliorating the metabolically adverse consequences of mild autonomous cortisol excess (MACS) due to adrenal adenomas (74) as affected patients are predominantly female (75).

Notably, quantification of AKR1C3 and HSD11B1 expression levels in subcutaneous and omental adipose tissue revealed a degree of interpersonal variation. Our computational simulations revealed that variations in the relative, as well as the absolute expression levels of HSD11B1 and AKR1C3 would result in individual differences in the amount of 11KT produced as well as the magnitude of change upon HSD11B1 inhibition. Indeed, this biological variation was reflected in both the *ex vivo* adipose tissue conversion data and the serum concentrations of 11KT before and after administration of AZD4017 in individuals with type 2 diabetes. Our simulations suggest that when considering 11KT biosynthesis, individuals with higher HSD11B1 expression relative to AKR1C3 would be less responsive to partial HSD11B1 inhibition. However, if HSD11B1 activity is completely abolished by inhibition then the amount of 11KT produced is dependent of the absolute levels of AKR1C3 expressed. Increased 11KT concentrations are therefore not solely dependent on the inhibition of HSD11B1, but also on the amount of AKR1C3 expressed. Notably, AKR1C3 expression correlates with BMI in subcutaneous, but not omental, adipose tissue and its expression is upregulated by insulin (14, 15, 17). Women with insulin resistance and obesity due to metabolic syndrome are therefore likely to have high expression levels of AKR1C3 (14, 15), which would result in greater increases in 11KT upon inhibition of HSD11B1. Thus, while such women may benefit from the local reduction of glucocorticoid activation by HSD11B1 inhibition in adipose tissue, they may be primed to generate more 11KT. Indeed, we noted that the expression of AKR1C3 was higher than that of HSD11B1 in adipose tissue obtained from women with obesity undergoing elective bariatric surgery. As a result, the conversion of 11KA4 yielded more 11KT and 11OHT combined (both requiring AKR1C3 activity) than 11OHA4. In subcutaneous adipose tissue, which had a lower HSD11B1:AKR1C3 ratio compared to omental adipose tissue, 11OHT was the most abundant product, demonstrating effective conversion of 11KA4 to 11KT by AKR1C3 and subsequent conversion of 11KT to 11OHT by HSD11B1. Similarly, we noted that in our clinical cohort, which consisted of individuals with type 2 diabetes (BMI mostly in the obese range), the mean circulating 11OHT concentration was ≥2-fold higher than previously reported for healthy individuals thus suggesting high peripheral AKR1C3 expression (3). Interestingly, a previous *ex vivo* study found that androgen exposure upregulates HSD11B1 expression in adipose tissue (76) emphasising that inactivation of 11-oxygenated androgens by HSD11B1 might be a protective mechanism, that no longer suffices once AKR1C3 is significantly upregulated, e.g. in the context of PCOS (15).

Increased 11KT concentrations following HSD11B1 inhibition are unlikely to be a concern in men as the metabolic effects of androgens are sexually dimorphic. In men, reduced androgen concentrations are associated with adverse metabolic consequences, including insulin resistance, type 2 diabetes and metabolic syndrome, and as such increased 11KT levels have the potential to be beneficial (43, 71, 77). Glucocorticoids are, however, also known to have sexually dimorphic effects, which are further complicated by crosstalk between glucocorticoids and androgens (78–80). Unfortunately, the vast majority of HSD11B1 inhibitor clinical studies did not report sex-specific outcomes for HSD11B1 inhibition (34–38). Larger sex-specific studies are therefore needed to fully assess the benefits of HSD11B1 inhibition in relation to metabolic conditions, while taking into account potential opposing androgenic effects in women, resulting from increased 11KT.

One recent study by Hardy *et al.* investigated the effects of HSD11B1 inhibition in women with obesity and Idiopathic Intracranial Hypertension (41). Twice daily administration of 400 mg AZD4017 for 12 weeks had no effect on fasting glucose, fasting insulin, insulin resistance or HbA1c, in agreement with our hypothesis that androgenic effects may counter beneficial metabolic effects of reduced glucocorticoid concentrations. Furthermore, this study found no effect of HSD11B1 inhibition on adipose or bone composition, but did report reduced cholesterol and increased HDL with HSD11B1 inhibition suggesting that women do exhibit some cardioprotective responses to HSD11B1 inhibition. Importantly, Hardy *et al*. also reported increased lean muscle mass with HSD11B1 inhibition, which correlated with increased testosterone and 11-oxygenated androgen precursors (11OHA4 and 11KA4) levels; however, they did not report 11KT concentrations.

Increased adrenal output of 11OHA4 is not unexpected with HSD11B1 inhibition, which leads to reduced negative feedback on the HPA-axis, increased ACTH levels and increased adrenal output of cortisol and adrenal androgen precursors (41). While we did not observe increased serum 11OHA4 concentrations in the current study, we cannot exclude that increased adrenal output of this precursor contributed to the observed increase in 11KT biosynthesis. In fact, this is likely the case as circulating 11OHA4 concentrations were not reduced despite inhibition of HSD11B1 and the resulting significant decrease in the 11OHA4/11KA4 ratio. Furthermore, the original clinical trial observed a significant increase in serum DHEAS, a marker of adrenal C19 steroid biosynthesis, thus confirming the stimulation of adrenal steroidogenesis (40). Despite the potential increase in adrenal 11OHA4 biosynthesis, our *in vitro*, *ex vivo* and *in silico* data unequivocally show that HSD11B1 modulates the peripheral biosynthesis of 11KT and that inhibition of HSD11B1 results in increased peripheral 11KT production irrespective of adrenal 11OHA4 output.

This study highlights that HSD11B1 inhibition may not be as effective as previously anticipated for the treatment of metabolic conditions in women due to the concomitant increase in the active androgen pool. However, the current data does not call into question the safety or efficacy of HSD11B1 inhibitors such as AZD4017, which has been shown to be well tolerated (39–41, 63, 81), but rather highlights that the condition targeted by the use of HSD11B1 inhibitors needs to be carefully considered. To date, AZD4017 has shown promise in the treatment of Idiopathic Intracranial Hypertension and wound healing in adults with type 2 diabetes, thus warranting further studies (40, 41). It may also be interesting to consider HSD11B1 inhibition in oophorectomized women. In this setting, the increased biosynthesis of 11KT due to HSD11B1 inhibition may be beneficial in offsetting the androgen deficiency due to oophorectomy, while reduced local glucocorticoid activation has the potential to protect against glucocorticoid induced osteoporosis, especially against the backdrop of reduced estrogen biosynthesis (82–84).

While the current study focusses on the regulation of 11-ketotestosterone biosynthesis by HSD11B1 in adipose tissue, HSD11B1 is also abundantly expressed in liver and as such the hepatic conversion of 11KA4 to 11OHA4 and 11KT to 11OHT may also regulate systemic 11KT levels, with inhibition of hepatic HSD11B1 by AZD4017 contributing to the observed increase in 11KT levels (18, 19).

In summary, our findings indicate that HSD11B1 plays an essential role in modulating the amount of 11KT produced by AKR1C3 expressed in adipose tissue. As a result, inhibition of HSD11B1 does not just reduce local glucocorticoid activation, but also alters the biosynthesis of the potent 11-oxygenated androgen 11KT. In women, the significant increase in 11KT concentrations may have metabolically opposing effects that attenuate the beneficial reduction of glucocorticoid activation in metabolic tissue. It is therefore imperative that future HSD11B1 inhibitor studies consider both the glucocorticoid and androgen pools and that sex-specific effects are systematically investigated and considered in the design of future studies.

## Data Availability

The data that support the findings of this study are available in the results and/or supplementary material of this article. The computational model is available in SBML format upon request.

## Conflict-of-interest

## Author contributions

KHS, WA and LS were responsible for study concept and design. LS and IO conducted experiments. JS performed all computational modelling. LS, IO, JS, LCG, AET, and KHS analysed data. AJC and RS provided human adipose tissue. AT, RA and AF provided serum and urine from a phase 2b pilot trial with AZD4017. AET and LCG conducted the analysis of clinical samples by LC-MS/MS and GC-MS, respectively. KHS, LS and IO drafted the manuscript. WA, AT and JS critically reviewed the manuscript. All authors reviewed and approved the final version of the manuscript. KS and WA obtained funding. These authors contributed equally: LS and IO. LS was selected as the first of the two authors as she was involved in the conceptualisation of the study.

