## Supplementary Material for "Inhibition of the glucocorticoid-activating enzyme 11β-hydroxysteroid dehydrogenase type 1 drives concurrent 11-oxygenated androgen excess"

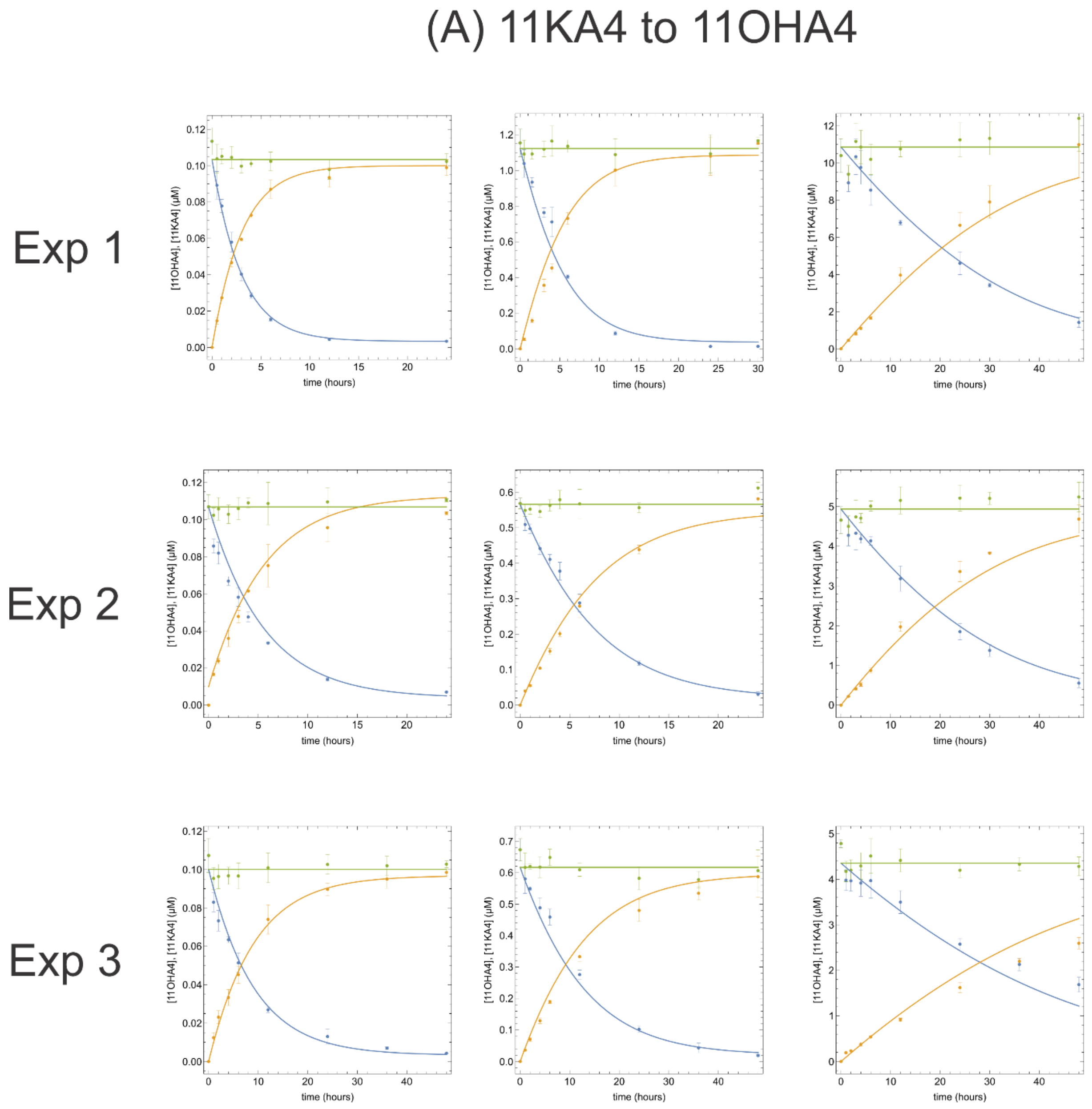

**Supplementary Material**
Schiffer et al. Inhibition of the glucocorticoid-activating enzyme 11β-hydroxysteroid dehydrogenase type 1 drives concurrent 11-oxygenated androgen excess

(B) 11KT to 11OHT

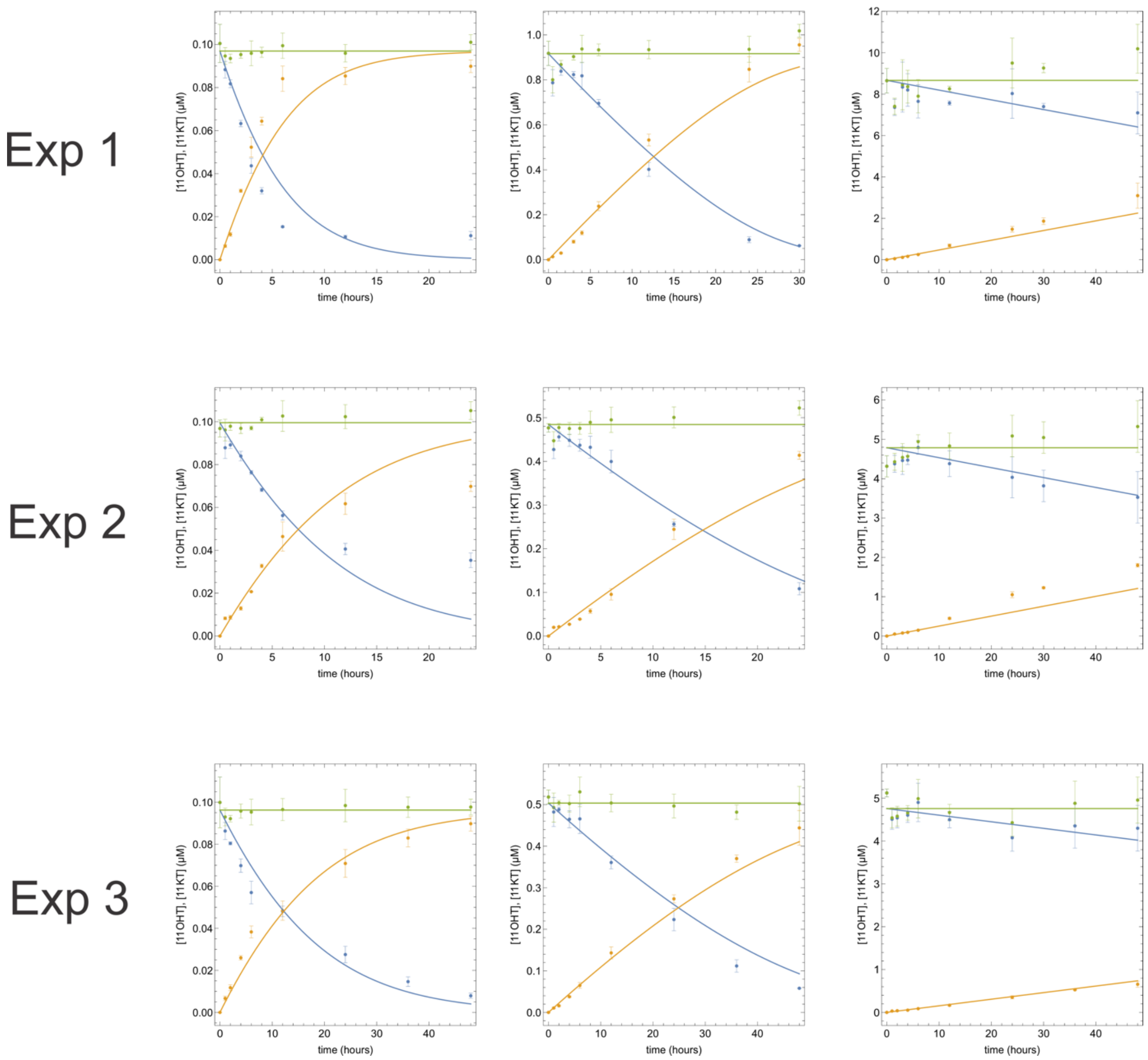

**Supplementary Material**

Schiffer et al. Inhibition of the glucocorticoid-activating enzyme 11β-hydroxysteroid dehydrogenase type 1 drives concurrent 11-oxygenated androgen excess

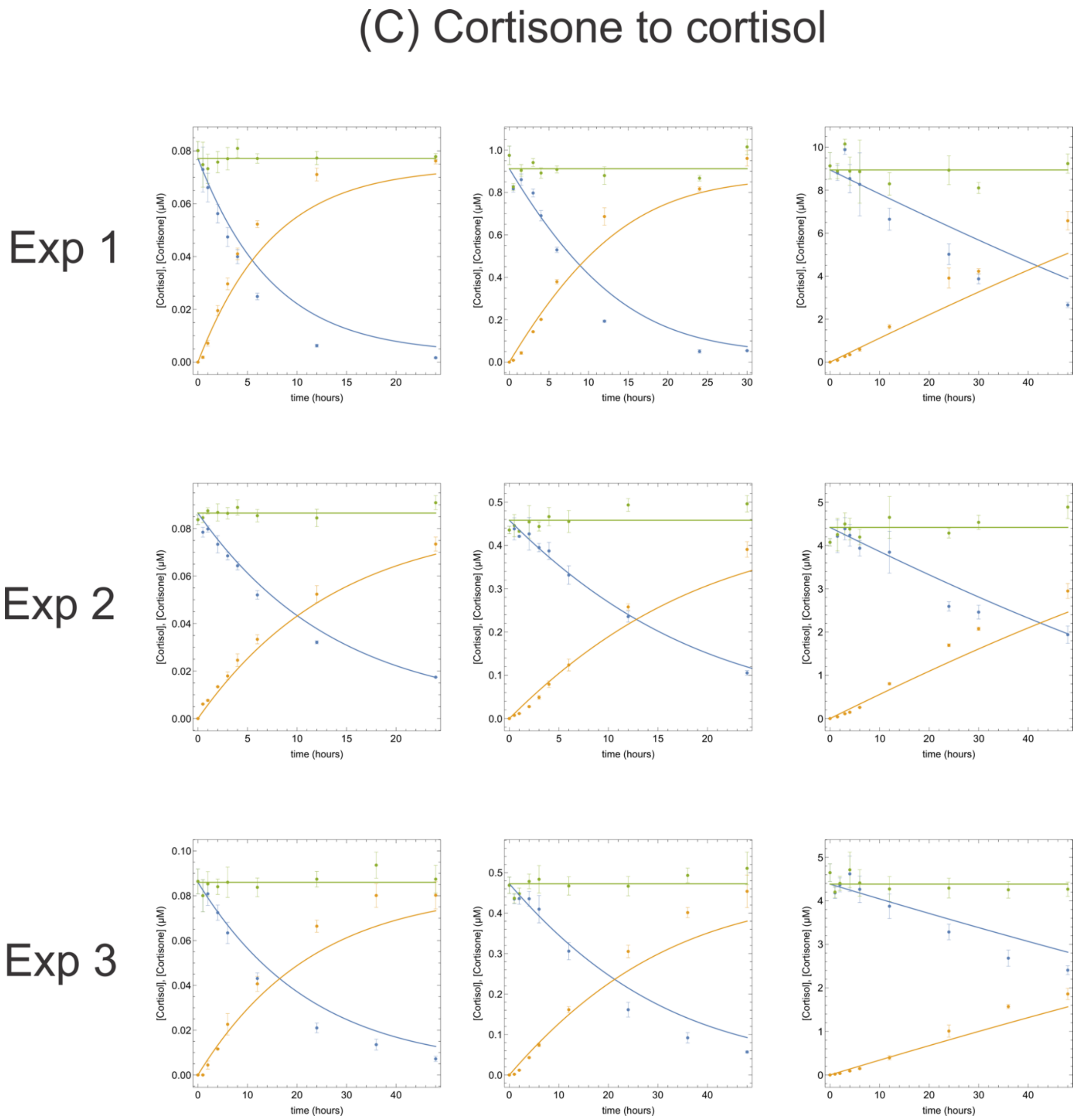


**Supplementary Figure 1: Time course experiments used to determine the apparent kinetic parameters ($K_{m,\ app}$ and $V_{max,\ app}$) of HSD11B1 towards 11KA4 (A), 11KT (B) and cortisone (C).** Experimental results are shown as the mean ±SD. The solid lines represent the model fits used to determine the apparent kinetic parameters. The blue lines are fits to the substrate concentration, the orange lines are fits to the product concentration, and the green lines represent the sum of the substate and product concentrations.

**Supplementary Material**
Schiffer et al. Inhibition of the glucocorticoid-activating enzyme 11β-hydroxysteroid dehydrogenase type 1 drives concurrent 11-oxygenated androgen excess

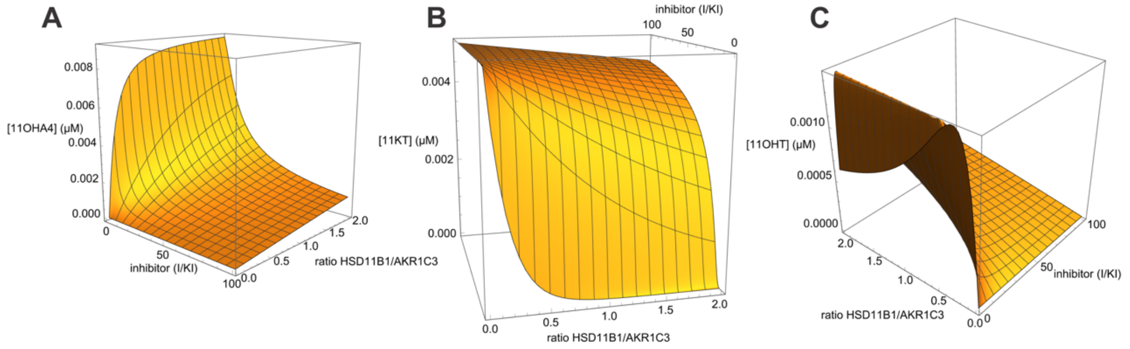


**Supplementary Figure 2**: **Computational model of the effect of varying HSD11B1:AKR1C3 ratios and different levels of HSD11B1 inhibition on the resulting concentrations of 11OHA4 (A), 11KT (B) and 11OHT (C).** The model varies that ratio of HSD11B1:AKR1C3 as in Figure 2 and estimates the effect of different levels of HSD11B1 inhibition (I/Ki = 0-100) on the conversion of 10 nM 11KA4 after 24-hours as in Figures 2 and 3.

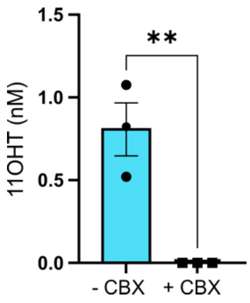


**Supplementary Figure 3**: **Inhibition of HSD11B1 by Carbenoxolone (CBX) results in decreased 11OHT production from 11KA4 when HSD11B1 and AKR1C3 are co-expressed (1:1 ratio).** Cells were treated with 100 nM cortisone and 10 nM 11KA4 with or without CBX (10 μM) for 24-hours after which steroid concentrations were determined by UHPLC-MS/MS. Results are shown as means ± SEM of three biological experiments performed in triplicate. P values were calculated using unpaired t-tests (**P<0.01)

**Supplementary Material**

Schiffer et al. Inhibition of the glucocorticoid-activating enzyme 11β-hydroxysteroid dehydrogenase type 1 drives concurrent 11-oxygenated androgen excess

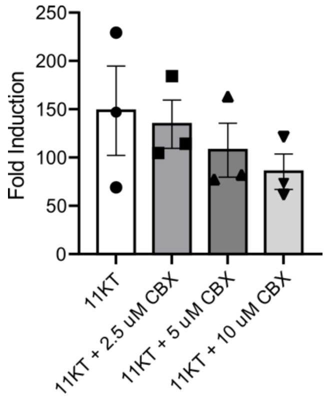


**Supplementary Figure 4**: **Carbenoxolone (CBX) antagonises the AR. HEK293 cells were transiently transfected with a plasmid expressing the androgen receptor (AR) and a luciferase reporter construct.** Cells were treated with 10 nM 11KT and increasing concentrations of CBX for 24-hours. Luciferase was subsequently assayed and is shown as the fold-induction relative to a vehicle control. All results are shown as the mean ± SEM of three biological experiments.

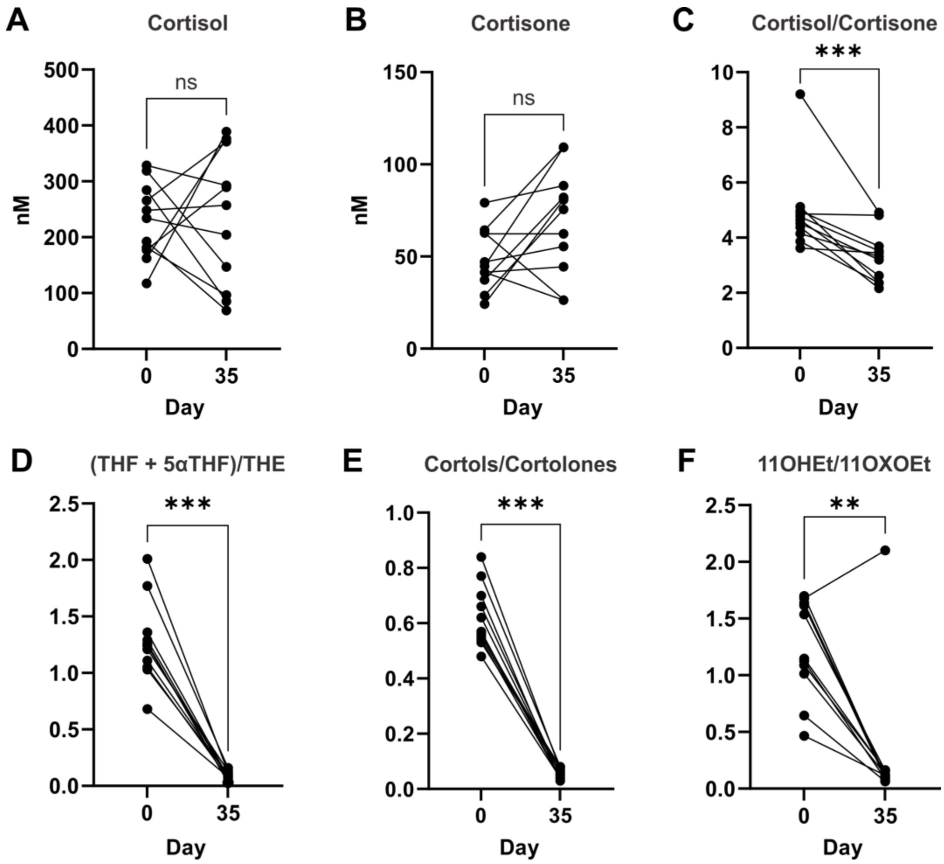


**Supplementary Figure 5**: **Inhibition of HSD11B1 by oral administration of AZD4017 (400 mg twice daily) for 35 days (n=11) changes the serum (A-C) and urinary (D-F) profile of glucocorticoids**. P values were calculated using Wilcoxon signed-rank tests (*P <0.05; **P<0.01; ***P<0.001).

**Supplementary Material**

Schiffer et al. Inhibition of the glucocorticoid-activating enzyme 11β-hydroxysteroid dehydrogenase type 1 drives concurrent 11-oxygenated androgen excess

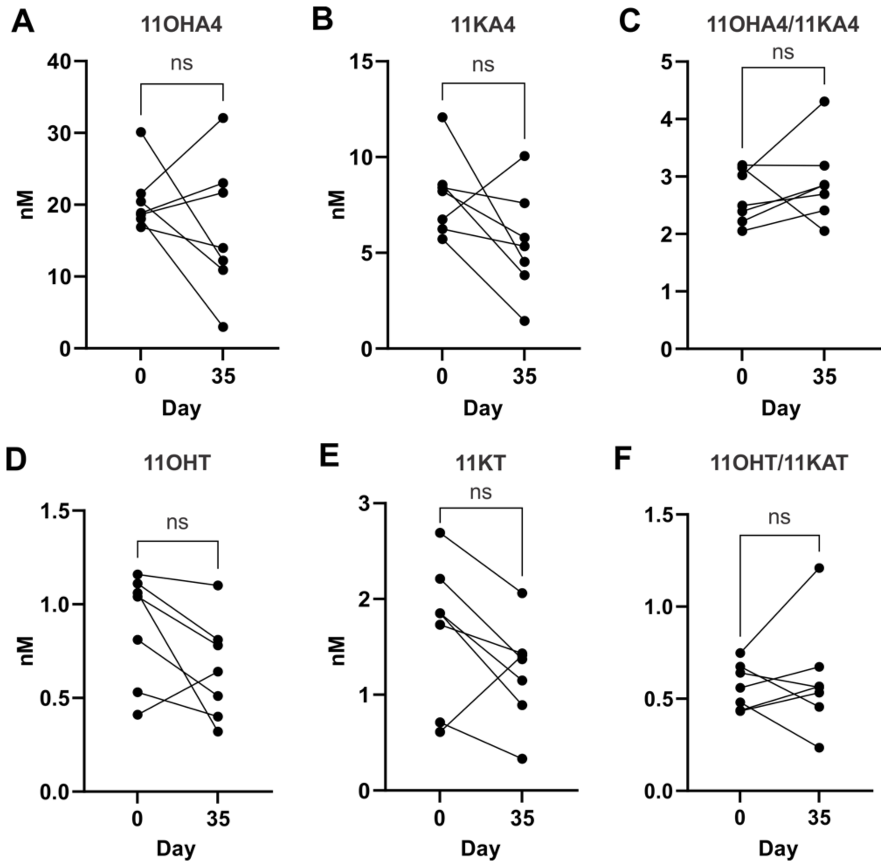


**Supplementary Figure 6: Serum concentrations of 11-oxygenated androgens for individuals receiving placebo.**

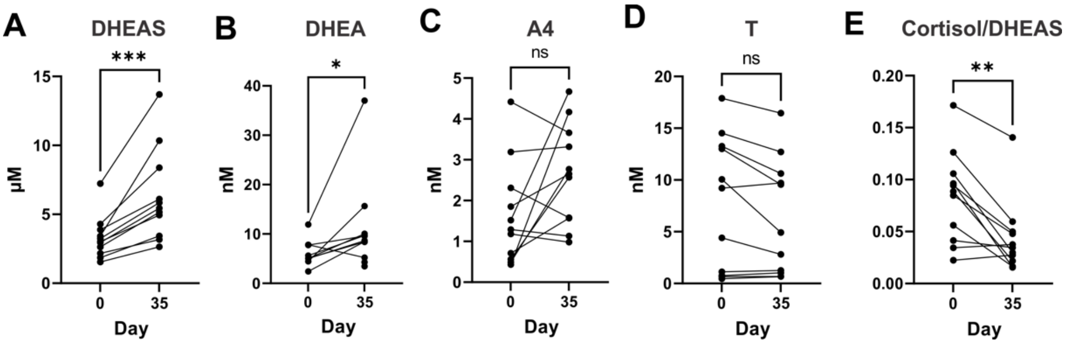


**Supplementary Figure 7**: **Effect of selective HSD11B1 inhibition on classic androgens (A-D) and the cortisol/DHEAS ratio (E) in circulation.** P values were calculated using Wilcoxon signed-rank tests (*P <0.05; **P<0.01; ***P<0.001).

**Supplementary Material**

Schiffer et al. Inhibition of the glucocorticoid-activating enzyme 11β-hydroxysteroid dehydrogenase type 1 drives concurrent 11-oxygenated androgen excess

**Supplementary Table 1: Correlation Analysis (Spearman's Rho, all participants) for change in glucocorticoid and 11-oxygenated androgen concentrations and ratios (Day 35-0) for all participants taking part in the study (twice daily oral administration of AZD4017 (400 mg) (n=11) or placebo (n=7)).**

| | Cortisol | Cortisone | Cortisol/ Cortisone | 11OHA4 | 11KA4 | 11OHA4 /11KA4 | 11KT | 11OHT | 11OHT/ 11KT |
|---|---|---|---|---|---|---|---|---|---|
| Cortisol | | **0.831[c]** | 0.439 | **0.903[c]** | **0.812[c]** | 0.238 | 0.379 | **0.722[c]** | 0.207 |
| Cortisone | **0.831[c]** | | -0.011 | **0.859[c]** | **0.932[c]** | -0.185 | **0.701[b]** | 0.420 | -0.203 |
| Cortisol/ Cortisone | 0.439 | -0.011 | | 0.208 | -0.038 | **0.893[c]** | -0.375 | 0.373 | **0.686[b]** |
| 11OHA4 | **0.903[c]** | **0.859[c]** | 0.208 | | **0.944[c]** | 0.047 | 0.452 | **0.603[b]** | **0.007[b]** |
| 11KA4 | **0.812[c]** | **0.932[c]** | -0.038 | **0.944[c]** | | -0.201 | **0.624[b]** | 0.439 | -0.199 |
| 11OHA4/ 11KA4 | 0.238 | -0.185 | **0.893[c]** | 0.047 | -0.201 | | -0.455 | 0.231 | **0.517[a]** |
| 11KT | 0.379 | **0.701[b]** | -0.375 | 0.452 | 0.624 | -0.455 | | -0.068 | **-0.624[b]** |
| 11OHT | **0.722[a]** | 0.420 | 0.373 | **0.603[b]** | **0.439[b]** | 0.231 | -0.068 | | **0.479[a]** |
| 11OHT/ 11KT | 0.207 | -0.203 | **0.686[b]** | 0.007 | -0.199 | **0.517[a]** | **-0.624$_b$** | **0.479[a]** | |

[a]P<0.05
[b]P<0.01
[c]P<0.001

**Model description**

The model for the combined simulation of 11HSDB1 and AKR1C3 activities is formulated in terms of ordinary differential equations:

$$\frac{d\,\mathrm{ka4}(t)}{d\,\mathrm{t}} = -v_{\mathrm{HSD\_ka4}} - v_{\mathrm{AKR\_ka4}}$$

$$\frac{d\,\mathrm{oha4}(t)}{d\,\mathrm{t}} = v_{\mathrm{HSD\_ka4}}$$

$$\frac{d\,\mathrm{kt}(t)}{d\,\mathrm{t}} = v_{\mathrm{AKR\_ka4}} - v_{\mathrm{HSD\_kt}}$$

$$\frac{d\,\mathrm{oht}(t)}{d\,\mathrm{t}} = v_{\mathrm{HSD\_kt}}$$

$$\frac{d\,\mathrm{cort}(t)}{d\,\mathrm{t}} = -v_{\mathrm{HSD\_cort}}$$

$$\frac{d\,\mathrm{cortsol}(t)}{d\,\mathrm{t}} = v_{\mathrm{HSD\_cort}}$$

With ka4=11KA4, oha4=11OHA4, kt=11KT, oht=11OHT, cort=cortisone and cortsol=cortisol. The rate equations used in the model for the two enzymes is dependent on the respective substrates that are converted, and the equations were parameterised by fitting them to progress curves for the individual substrates (**Supplementary Figure 1**). Note that HSD11B1 catalyses three reactions and that the substrates/products of each of these reactions can in principle act as competitive inhibitors for the other reactions. In the rate equations displayed below, all effectors that were observed to affect the reaction rates are indicated, (in case an effector is not present in a specific incubation its concentration is simply set to 0).

**Supplementary Material**

Schiffer et al. Inhibition of the glucocorticoid-activating enzyme 11β-hydroxysteroid dehydrogenase type 1 drives concurrent 11-oxygenated androgen excess

$$v_{\mathrm{HSD\_ka4}} = \frac{\frac{V_{\mathrm{Mf\_HSDka4}}\cdot[ka4(t)]}{K_{\mathrm{M\_HSDka4}}} - \frac{V_{\mathrm{Mr\_HSDka4}}\cdot[oha4(t)]}{K_{\mathrm{M\_HSDoha4}}}}{1 + \frac{[ka4(t)]}{K_{\mathrm{M\_HSDka4}}} + \frac{[kt(t)]}{K_{\mathrm{M\_HSDkt}}} + \frac{[oha4(t)]}{K_{\mathrm{M\_HSDoha4}}} + \frac{[cort(t)]}{K_{\mathrm{M\_HSDcort}}}}$$

$$v_{\mathrm{HSD\_kt}} = \frac{\frac{V_{\mathrm{Mf\_HSDkt}}\cdot[kt(t)]}{K_{\mathrm{M\_HSDkt}}}}{1 + \frac{[ka4(t)]}{K_{\mathrm{M\_HSDka4}}} + \frac{[kt(t)]}{K_{\mathrm{M\_HSDkt}}} + \frac{[oha4(t)]}{K_{\mathrm{M\_HSDoha4}}} + \frac{[cort(t)]}{K_{\mathrm{M\_HSDcort}}}}$$

$$v_{\mathrm{HSD\_cort}} = \frac{\frac{V_{\mathrm{Mf\_HSDcort}}\cdot[cort(t)]}{K_{\mathrm{M\_HSDcort}}} - V_{\mathrm{Mr\_HSDcort}}\cdot[cortsol(t)]}{1 + \frac{[ka4(t)]}{K_{\mathrm{M\_HSDka4}}} + \frac{[kt(t)]}{K_{\mathrm{M\_HSDkt}}} + \frac{[oha4(t)]}{K_{\mathrm{M\_HSDoha4}}} + \frac{[cort(t)]}{K_{\mathrm{M\_HSDcort}}}}$$

$$v_{\mathrm{AKR\_ka4}} = \frac{\frac{V_{\mathrm{M\_AKRka4}}\cdot[ka4(t)]}{K_{\mathrm{M\_AKRka4}}}}{1 + \frac{[ka4(t)]}{K_{\mathrm{M\_AKRka4}}}}$$

**Supplementary Table 2: Kinetic constants for HSD11B1 and AKR1C3*.**

| | | |
|---|---|---|
| $V_{Mf\_HSDka4}$ | 0.35 | (µM/h) |
| $V_{Mr\_HSDka4}$ | 0.03 | (µM/h) |
| $V_{Mf\_HSDkt}$ | 0.05 | (µM/h) |
| $V_{Mf\_HSDcort}$ | 0.12 | (µM/h) |
| $V_{Mr\_HSDcort}$ | 0.01 | (1/h) |
| $K_{M\_HSDka4}$ | 1.0 | (µM) |
| $K_{M\_HSDkt}$ | 0.21 | (µM) |
| $K_{M\_HSDoha4}$ | 2.46 | (µM) |
| $K_{M\_HSDcort}$ | 0.9 | (µM) |

| | | |
|---|---|---|
| $V_{M\_AKRka4}$ | 0.27 | (µM/h) |
| $K_{M\_AKRka4}$ | 1.87 | (µM) |

*All $V_M$ values are apparent values and dependent on transfection efficiency that differ between experiments. We included the conversion of 100 nM 11KA4 to 11OHA4 in all experiments and used the initial rate to normalise the transfection efficiency, which was set to 1 for the values reported above.
